# Directing polymorph specific calcium carbonate formation with de novo protein templates

**DOI:** 10.1101/2023.06.09.544362

**Authors:** Fatima A. Davila-Hernandez, Biao Jin, Harley Pyles, Shuai Zhang, Zheming Wang, Timothy F. Huddy, Chun-Long Chen, James J. De Yoreo, David Baker

## Abstract

Biomolecules modulate inorganic crystallization to generate hierarchically structured biominerals^1–5^, but the atomic structure of the organic-inorganic interfaces that regulate mineralization remain unknown^6–8^. We hypothesized that heterogeneous nucleation of calcium carbonate could be achieved by a structured flat molecular template that pre-organizes calcium ions on its surface. To test this hypothesis, we designed helical repeat proteins (DHRs) displaying regularly spaced carboxylate arrays on their surfaces and found that both protein monomers and protein-Ca^2+^ assemblies directly nucleate nano-calcite with non-natural (110) or (202) faces while vaterite, which forms first absent the proteins, is bypassed. The nanocrystals then assemble by oriented attachment into calcite mesocrystals. We find further that nanocrystal size and polymorph can be tuned by varying the length and surface chemistry of the designed protein templates. Thus, bio-mineralization can be programmed using de novo protein design, providing a route to next-generation hybrid materials.

**One sentence summary:** De novo designed protein templates promote nucleation of nano-calcite and direct its growth by oriented particle attachment.

## Introduction

Organisms generate biominerals such as nacre, shell, bone, and tooth^9,10^ by controlling the nucleation, growth, and assembly of inorganic crystals^1–5^. The sequences of native biomineral-associated proteins and their effects on mineralization are well characterized for some systems^3,11–19^ but the three-dimensional structures of the protein-mineral interfaces are largely unknown. For example, natural proteins nucleate specific calcium carbonate (CaCO_3_) polymorphs^13,14,20–22^, but none have known stable tertiary structures and many are intrinsically disordered or insoluble^6–8^. Additives including small organic molecules^23–25^, polymers^26^, amino acids^27^, and peptides^28,29^ affect CaCO_3_ crystallization, but have not been shown to template nucleation. CaCO_3_ growth modification and localization of nucleation has been explored with short unstructured CaCO_3_ binding peptides genetically fused to proteins immobilized on a surface and with post-translational modified proteins^30,31^, but in such systems it is difficult to structurally define the protein-mineral interface. Stereochemical matched self-assembled monolayers (SAM)^32^ modulate the nucleation of CaCO_3_, suggesting that geometric lattice matching may provide a general route to controlling mineralization^1,3,33,34^. In further support of this concept, the structures of ice binding proteins contain surfaces with an epitaxial-like lattice matching to the ice lattice^35^, which enables modulation of ice formation by binding ice nuclei through preorganized ice-like waters.

Guided by the ice binding protein example, we hypothesized that designed proteins with flat surfaces displaying functional groups lattice matched to a mineral of interest could be used to modulate mineralization by promoting ion association in a pattern consistent with the mineral lattice, thus directly reducing the free energy of formation of the critical nucleus (Fig. 1a). Advances in protein design now enable the creation of new proteins with precisely specified structures, and in previous work we showed that lattice matched interactions between arrays of carboxylate side chains on a designed protein surface and preformed mica crystals direct formation of precise protein-mica assemblies ^36^. As a first test of the potential of designed proteins for directly nucleating inorganic mineralization, we chose CaCO_3_ as a model system due to its numerous polymorphs, well documented nucleation behavior with and without proteins^37^, and prevalence in natural biominerals, and sought to design proteins with structures suitable for nucleating crystalline CaCO_3_ (Fig. 1a). We aimed to generate flat repetitive surface arrays of calcium-coordinating carboxylate groups (Fig. 1b) that could bind to and stabilize CaCO_3_ nuclei (Fig. 1c). To enable exploration of the contribution of the size of the protein-mineral interface while keeping the chemical composition and geometry otherwise constant, we employed Designed Helical Repeat (DHR; ^38^) proteins that can be shrunk or extended simply by changing the number of repeating units. These proteins have >10 nm^2^ flat surfaces with comparable or better structural definition and superior tunability relative to SAMs, and within genetically encoded soluble molecules that can be readily reprogrammed.

**Fig. 1.**
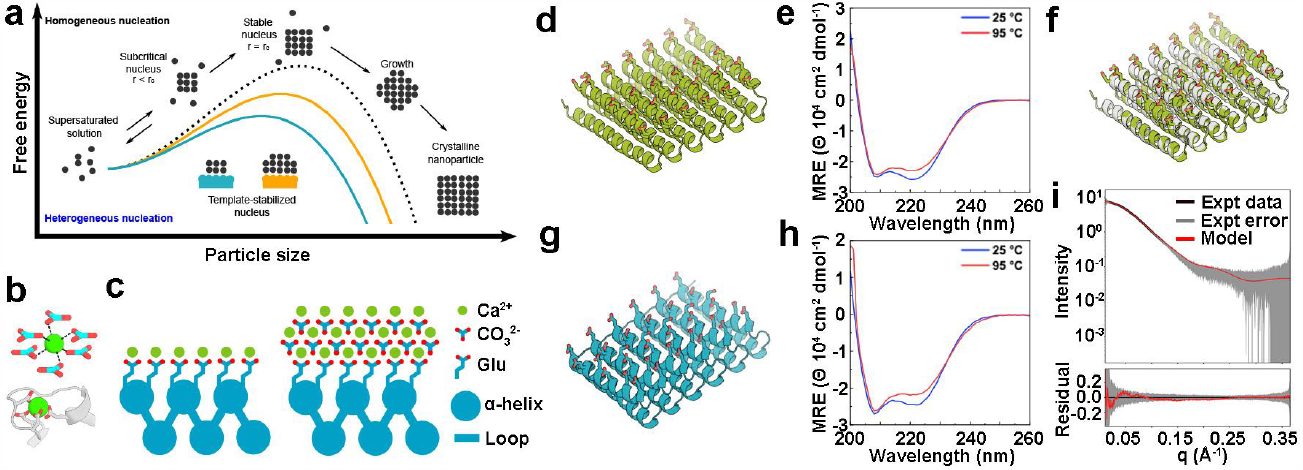
Design principles for the heterogeneous nucleation of CaCO3 guided by a protein template. (a) The presence of a template reduces the free energy barrier of nucleation and the critical radius of the nuclei. (b) Coordination of calcium (top) by carbonate ions within a unit cell of calcite, and (bottom) by carboxylate containing glutamate and aspartate residues in the structure of calmodulin (PDB ID: 1A29, residues 18-33). (c) Tessellating binding moieties across repeated α-helices within a designed protein capable of pre-organizing bound calcium ions. (d-i) Biophysical characterization of the designed proteins. Design models for (d) FD15 and (g) FD31. (e,h) Their respective circular dichroism scans showing mean residue ellipticity (M.R.E.) from 200 to 260 nm at 25 °C and 95 °C. (f) The crystal structure of FD15 in gray overlaid on the design model in green (solved at 3 Å resolution, RMSD of 0.45 Å). (i) Experimental and model SAXS profiles for FD31, scattering vector (q, from 0 to 0.35 Å-1) vs. intensity.

## Results

Flat repeat helix-turn-helix-turn protein backbones were designed with either constrained monte-carlo fragment assembly with RosettaRemodel ^39^, or by placing α-helices in a plane and then connecting them with additional secondary structural elements^40^. In contrast to typical designed protein surfaces containing a mix of positive, negative, and non-charged hydrophilic residues to promote solubility, and to hydrophobic protein surfaces designed to bind other proteins, we designed surfaces consisting entirely of carboxylate side chains spaced at regular intervals to mimic the carbonate sublattice on various crystallographic planes of CaCO_3_ crystals (supplemental methods).

We selected 40 designs for experimental characterization and obtained synthetic genes encoding them (Sup Table 1). Eight of these expressed at high levels in E. coli and were highly soluble. Two designs, FD15 and FD31, were selected for detailed characterization. Size exclusion chromatography (SEC) showed that both proteins were monodisperse in solution with molecular weights expected for the protein monomer (Fig. S1a,c). Circular dichroism (CD) experiments showed the expected alpha-helical structure, and the proteins were found to be hyperstable, remaining folded at temperatures up to 95 ºC (Fig. 1e,h). Small angle X-ray scattering (SAXS) profiles of FD31 were consistent with profiles computed for the design model (Fig. 1i). The crystal structure of FD15 was solved at 3 Å resolution and matched the design model with an RMSD of 0.45 Å (Fig. 1f). This perfectly straight repeat protein possesses a 8.7 Å spacing between helices in adjacent repeat subunits. In order to allow for protein-protein assembly, as in some native biomineralization proteins^41,42^, we also designed an array of carboxylate side chains onto the surface of DHR49, a previously designed flat DHR ^38^, and removed the capping elements on the terminal repeats to produce head-to-tail protein-protein interfaces. This protein (DHR49-Neg) is alpha-helical and thermostable (Fig. S1d) and AFM experiments show it indeed assembles end-to-end to form single protein diameter fibers at the mica-water interface (Fig. S1e).

We first investigated the effect of the designed proteins on CaCO_3_ mineralization using in-situ liquid-phase attenuated total reflectance-Fourier transform infrared spectroscopy (ATR-FTIR) and time-dependent ex-situ transmission electron microscopy (TEM) (Fig. 2). We added 1 uM of each designed protein or a BSA control to a mineralization solution of 5 mM CaCl_2_ and 5 mM NaHCO_3_, which is supersaturated for both vaterite and calcite but undersaturated for amorphous calcium carbonate (ACC). Vaterite appeared as the first solid phase in the protein-free mineralization solution and in the presence of bovine serum albumin (BSA), which has the large negative charge of the DHR proteins but lacks both the periodic array of carboxylates and flat interface (Fig. 2c,l). The vaterite crystals grew to become hundreds of nm diameter spheroidal particles. In contrast, in the presence of DHR49-Neg predominantly calcite nanocrystals ∼5 nm in size were initially observed (Fig. 2e,f). FD31 was also highly effective at promoting the direct nucleation of nanocrystalline calcite (Fig. 2h,i) as indicated by selected area electron diffraction (SAED) (Fig. S2a), cryo-TEM (Fig. S2b), and high-resolution (HR) TEM (Fig. S2c), and liquid-phase (LP)-TEM images (Fig. S2d). In contrast, FD15 did not alter the crystallization pathway from that seen with the no protein and BSA controls (Fig. 2k). Given that FD15, FD31, and DHR49-Neg all have flat surfaces with regularly arrayed carboxylate groups, these differences likely stem from differences in repeat spacing (Fig. S3) and detailed surface chemistry (Table S2).

**Fig. 2.**
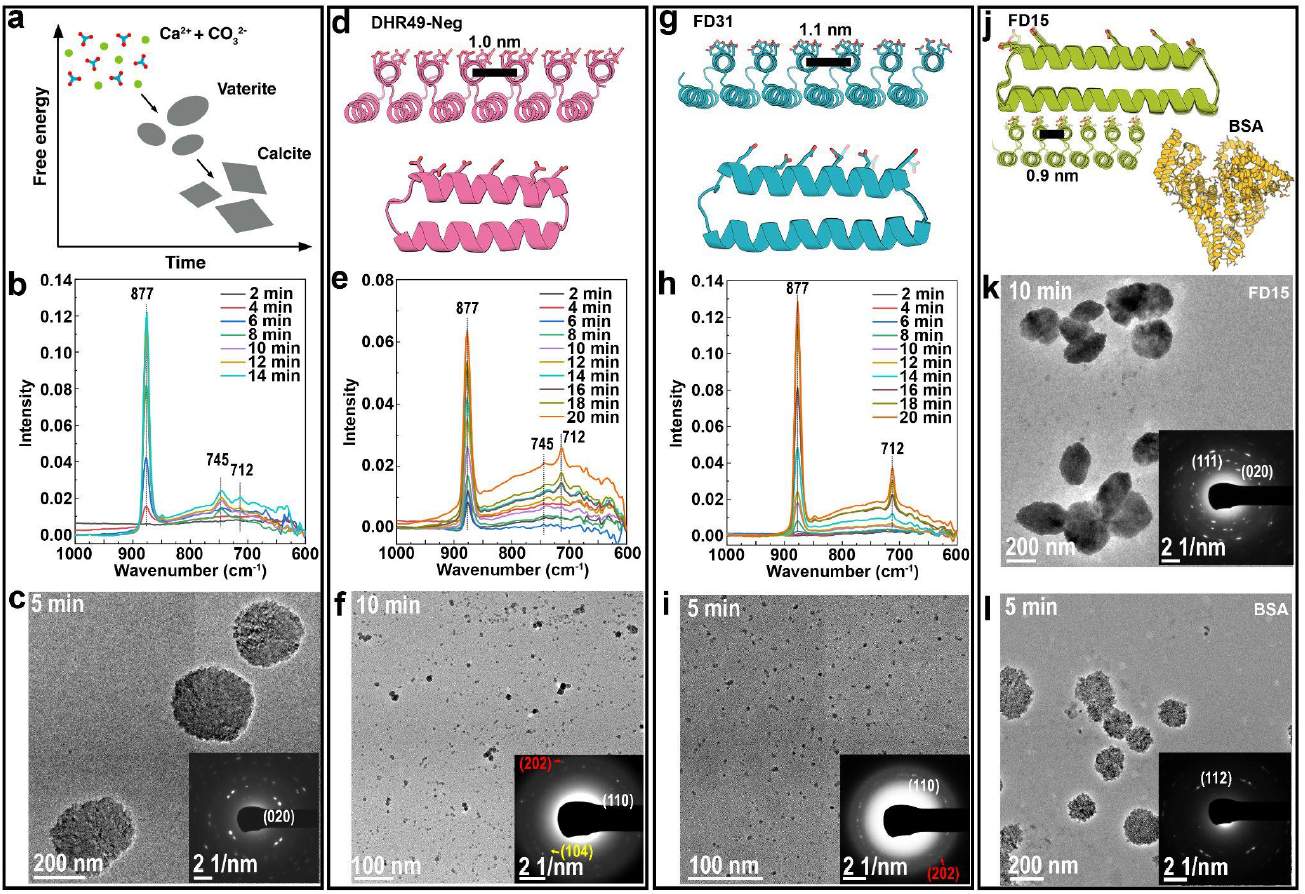
Designed proteins modulate CaCO3 crystallization. The CaCO3 crystallization process in supersaturated solutions containing 5 mM CaCl_2_ and 5 mM NaHCO_3_ was monitored in the absence and presence of different designed proteins. (a) Schematic showing the nucleation and transformation of calcium carbonate in the absence of additives under the conditions used here. First, vaterite forms followed by transformation to micron-sized calcite ^43^. (b) In situ ATR-FTIR and (c) TEM of CaCO3 crystallization in the absence of protein showing formation of vaterite. (d-l) Impact of 1 μM designed proteins on mineralization. Protein design models (d,g) and corresponding in situ ATR-FTIR (e,h) showing formation of predominantly calcite in the presence of DHR49-Neg, and exclusively calcite in the presence of FD31. Representative TEM images confirm the formation of (c) vaterite in the absence of the proteins, (f) primarily calcite crystals with DHR49-Neg, and (i) calcite nanocrystals in the presence of FD31. In contrast, in the presence of (j) FD15 or a BSA control (j) only vaterite is formed (k,l).

To investigate the mechanism underlying these nucleation effects we focused on FD31, which effectively promoted direct formation of nano-calcite. Supersaturated solutions containing 1 μM protein were sealed into a liquid cell for LP-TEM observations. Two nucleation pathways were observed. In the first, calcite nanocrystals nucleated throughout the solution (Fig. 3a; Movie S1), reaching a diameter of ∼7 nm and exhibiting a number density comparable to that of the protein monomers (∼1.1E15 *vs* ∼6.5E14; See supporting discussion for details). Increasing the protein concentration from 0.5 μM to 1 μM and 2 μM led to an increase in the number density of nano-calcite particles (Fig. S5a-c). Due to their low contrast and small size, individual protein monomers are not observable with LP-TEM, however AFM indicates their presence in solution (Fig. S4e). The calcite nanoparticles were also observed when imaging was performed at lower magnification and were already present when the electron beam was moved to neighboring regions (Fig. 3b), indicating that electron beam effects were not driving the process (See supporting text for more details). FD31 is 8 nm by 5 nm by 3 nm in size, which is similar to the 1∼5 nm critical nucleus size of CaCO_3_ ^44,45^. The similarity in size and number density of the calcite nanocrystals and protein monomers suggests nucleation is driven by individual proteins.

**Fig. 3.**
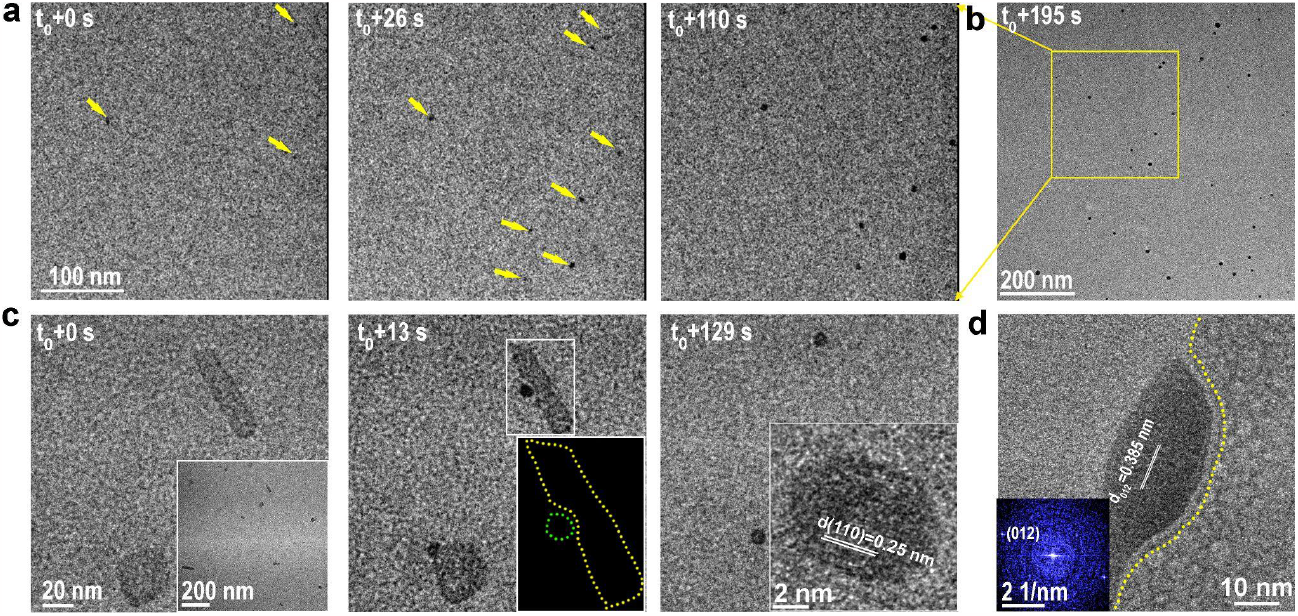
FD31 mediated nucleation of calcite nanocrystals. (a) Sequential LP-TEM images show the nucleation of eight CaCO_3_ nanoparticles in a precursor solution containing FD31 protein. (b) Low-mag LP-TEM image shows more nucleated nanoparticles. (c) Sequential LP-TEM images show the initial nucleation and growth of two CaCO_3_ nanoparticles around the sheet-like templates after FD31 was preincubated with CaCl_2_ for 5 minutes before addition of NaHCO_3_. Inset at t_0_ + 0 s displays multiple template formation at an early stage. A schematic (Inset at t_0_+13 s) shows the disconnected interface with a width of ∼1 nm between the as-formed particle and the substrate. At t_0_+129 s, the protein assembly dissolves, likely due to depletion of Ca, and the protein disperses into solution. Inset at t_0_+129 s shows LP-HR-TEM image of newly formed particle with the (110) lattice spacing of calcite. (d) In situ LP-HR-TEM image reveals the interface structure between calcite and the template. The FFT image in the inset shows the nucleated particle is calcite. As in Fig. 2, FD31 protein is at 1 μM and CaCl_2_ and NaHCO_3_ at 5 mM.

We observed a second nucleation pathway driven by sheet-like assemblies when FD31 was incubated with Ca^2+^ prior to addition of NaHCO_3_ (Fig. 3c; Movie S2). Preincubation of 2 μM FD31 with 10 mM Ca^2+^ in the absence of HCO_3_^-^, led to the formation of supramolecular assemblies (inset in Fig. 3c). Upon addition of an equal volume of 10 mM NaHCO_3_, after 5 mins we observed the nucleation of nanoparticles adjacent to the protein-Ca^2+^ supramolecular assemblies (Fig. 3c, Movie S2, Fig. S4j) identified as calcite by in situ high resolution (HR)-TEM (Inset in Fig. 3c). As the calcite particles continued to grow, the protein-Ca^2+^ assemblies dissolved (Fig. 3c). Incubation of FD31 with CaCl_2_ for 30 mins to 24 hours led to coordination of Ca by carboxylate groups (Fig. S4a,b) accompanied by formation of the protein-Ca^2+^ assemblies, which were found to be crystalline (Fig. S4d-i, Fig. S6). The protein-Ca^2+^ assemblies likely dissolve as calcite particles grow because the available Ca^2+^ in their vicinity is depleted.

At favorable viewing angles, a ∼1 nm separation was observed between the protein-Ca^2+^ assemblies and the nascent calcite nanocrystals (Fig. 3c,d). This is consistent with the ∼1 nm thickness of hydration layering on calcite ^46^ and suggests nucleation is an interface-driven process, as observed in other mineral systems in which surface ligands present carboxyl rich interfaces^47,48^. The results suggest that the flat periodic array of acidic residues on the surface of the proteins direct the absorption and organization of Ca^2+^ ions and water molecules in the hydration layer into an arrangement consistent with the structure of calcite. Alternatively, the interfacial region may alter the activities of the CaCO_3_ phases to modulate the supersaturation with respect to calcite vs vaterite.

We next followed the growth of FD31 nucleated calcite nanocrystals over time using LP-TEM. During the ∼5 min following nucleation, individual nuclei assembled into ∼10-20 nm particles through multiple attachment events, rather than individually growing in size (Fig. 4a, Movie S3). Ex-situ HR-TEM shows the larger agglomerated particles are surrounded by smaller particles (Fig. 4b-c), and the assemblies have continuous crystal lattices (Fig. 4d). Similar agglomerates were also observed after growth with DHR49-Neg at this time point (Fig. S7). Based on these observations we conclude that FD31 and DHR49-Neg stabilize calcite crystals at ∼5 nm size and that oriented particle attachment^4^ is the dominant pathway for calcite growth in their presence.

**Fig. 4.**
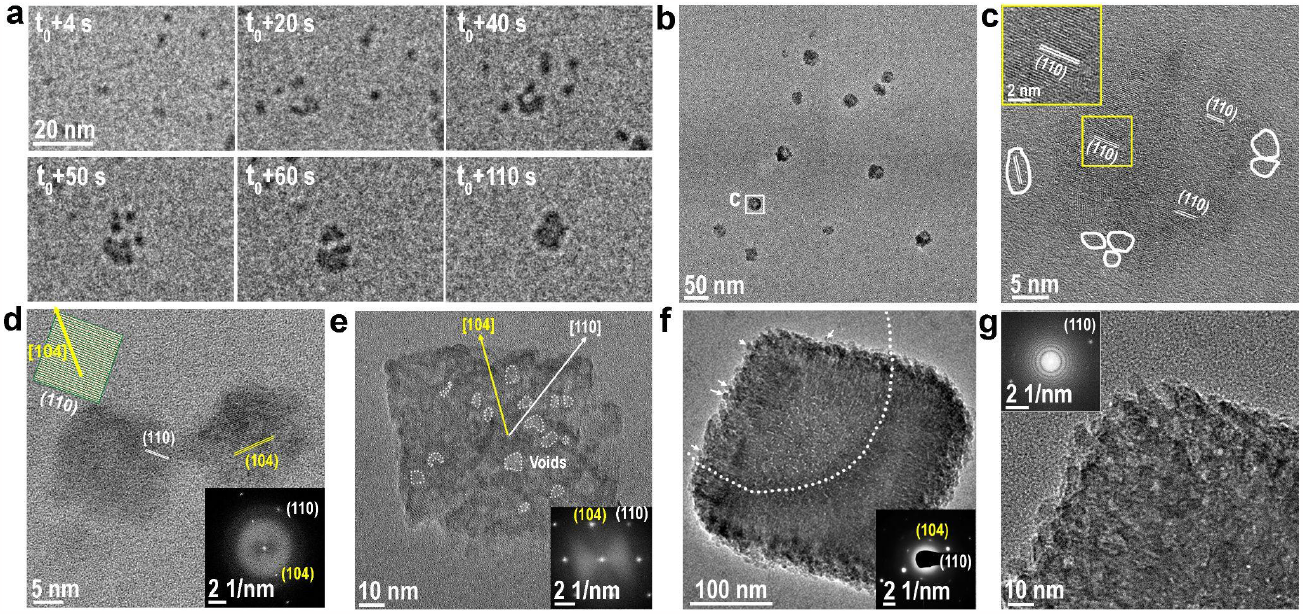
Calcite nanocrystal growth and assembly in the presence of FD31. (a) Sequential LP-TEM image sequence shows the attachment of multiple calcite nanocrystals into a larger one. (b) Ex situ TEM image showing calcite grown through particle attachment. (c) HR-TEM image showing the aggregated crystal with (110) lattice orientation. White frames mark several individual calcite nanocrystals around the larger particle. (d) HR-TEM image shows additional calcite nanocrystals with a size of ∼20 nm and (110) side faces. (e) TEM image shows a ∼100 nm rhombohedron-like calcite. The inset FTT shows the crystal orientation and single crystallinity. (f) TEM image showing rhombohedral calcite with a size of ∼300 nm. The embedded SAED image demonstrates its single crystalline nature. Arrows mark several calcite nanocrystals on the surface. (g) HR-TEM and corresponding FFT images in the inset confirm the single crystalline nature and that the rough surface is composed of attached nanoparticles.

The ∼10-20 nm agglomerated calcite crystals formed with FD31 and DHR49-Neg also express different preferred facet orientations, (110) for FD31 (Fig. 4b-d) and (202) for DHR49-Neg (Fig. S7c,d), both of which are distinct from the natural (104) facets of calcite grown in protein-free solutions. The differences between facet orientations for the two proteins likely reflect differences in their structure and chemistry (Highlighted in Table S2). First, while both are composed of repeating alpha helices, these are spaced at different distances, potentially leading to different extents of epitaxially matching for different calcite surfaces (see supplementary discussion, Fig. S3). Second, although the two proteins have similar net-charges and number of carboxylate groups, they contain different ratios of aspartate and glutamate residues, which present carboxylate groups in different orientations that result in distinct stereochemical alignments relative to carbonate ions in the calcite surface (see supplementary discussion, Fig. S8).

At 20 min in FD31 solutions we observed ∼100 nm calcite crystals with a pseudo-rhombohedral morphology (Fig. 4e), their habit resembling the thermodynamically stable (104) calcite rhombohedron, but with terraced corners and low contrast features that suggest a discontinuous lattice separated by voids, perhaps created by the inclusion of proteins. At 30-40 mins, the calcite crystals grew to ∼300 nm to ∼800 nm (Fig. 4f,g; Fig. S9 a-b), and retained a similar morphology with rough surfaces, internal voids, and multiple 4-8 nm particles on their surfaces (Fig. S9 c-h), suggesting growth through repeated attachment of the nanocrystals. This conclusion is supported by LP-TEM data (Fig. S10e). When the FD31 concentration was increased fourfold, the calcite crystals were elongated along the c-axis ([001] direction) (Fig. S11i-l). DHR49-Neg also formed calcite rhombohedra of similar size and habit (Fig. S7g). Thus, in the presence of the proteins, the 4-8 nm primary calcite nanocrystals assemble through oriented attachment to ultimately form micron-scale mesocrystalline rhombohedral calcite.

We next investigated what specific features of FD31 contribute to its potent calcite nucleation activity by generating versions of the protein with varied structural and chemical features. The effect of template length (the extent of the carboxylate arrays) was investigated by generating versions of the protein with different numbers of repeat subunits. FD31 has 6 repeat units and a 5 × 8 nm interface containing 36 carboxylate groups. We produced a 3 repeat version with a 5 × 5 nm interface containing 18 carboxylate groups and a 9 repeat version with a 5 × 11 interface containing 54 carboxylate groups. The 3, 6, and 9 repeat templates all drive the formation of nano-calcite, but the 3 repeat version leads to larger nanoparticles (Fig. 5a-c), likely due to reduced ability to lower the interfacial energy or weaker binding to calcite surfaces following nucleation (see below for discussion). Nucleation driven by monomers of FD31-Rep9 was also observed by liquid-phase TEM (Fig. S12).

**Fig. 5.**
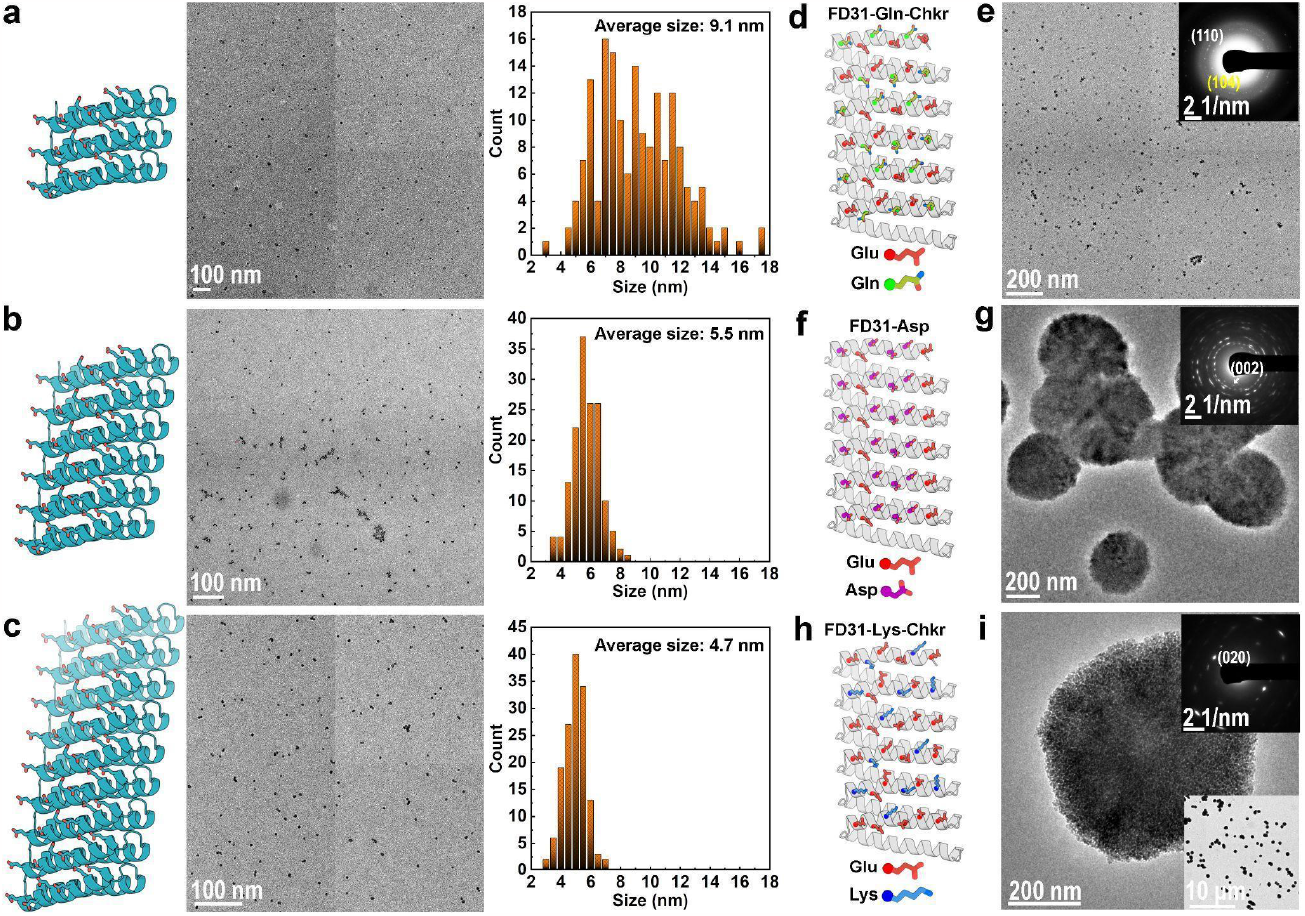
Tuning calcite nucleation by varying FD31 length and surface chemistry. (a-c) TEM and the size distribution of particles formed in the presence of (a) FD31-Rep3 with 3 repeats, (b) FD31 with 6 repeats, and (c) FD31-Rep9 with 9 repeats. (d-i) TEM and SAED show the formation of (e) calcite-dominant nanocrystals in the presence of FD31-Gln-Checker (d); the formation of (g) vaterite as well as (Fig. S13c,d) calcite in the presence of (f) FD31-Asp; and the formation of a (i) vaterite dominant phase in the presence of (h) FD31-Lys-Checker.

To investigate the effect of surface chemistry on calcite nucleation activity, we modified the surface chemistry of FD31 in three ways. We either substituted half of the glutamate residues to nearly isosteric but non-charged glutamine residues (FD31-Gln-Checker), replaced 24 out of 36 glutamate residues with shorter aspartic acid residues (FD31-Asp), or mutated 13 out of 36 of the glutamate to basic lysine residues (FD31-Lys-Checker)(Fig. 5d,f,h). FD31 (containing an all glutamate interface) and FD31-Gln-Checker generated calcite-dominant nanocrystals (Fig. 2g,h,i, Fig. 5e, Fig. S13a-d), both vaterite and calcite phases appeared in the presence of FD31-Asp (Fig. 5g, Fig. S13c-d), and FD31-Lys-Checker produced primarily vaterite (Fig. 5i, S13i-l). Thus the structure and chemistry of the side chain arrays in our designed protein directly influence the templating of calcite nucleation.

## Discussion

Our designed protein templates promote the direct formation of 5-9 nm calcite nanocrystals through both monomer- and protein-Ca^2+^ assembly-driven nucleation, bypassing the formation of vaterite microcrystals that occurs in their absence or with other negative design controls. The latter pathway resembles the multistep process in bone and dentin calcification where Ca^2+^ induced assembly of an acidic matrix protein proceeds formation of apatite^49^. The proteins then inhibit further growth of the crystals directly from solution, stabilize non-natural faces, and switch the crystallization pathway to nonclassical oriented attachment, ultimately producing micron-scale rhombohedral single crystals with terraced edges, abundant low contrast inclusions and morphologies that vary with protein design and concentration. The direct appearance of calcite strongly suggests that the distributions of carboxylic side chains bias the configuration of the ions at the protein-solution interface towards that of the calcite lattice, but determination of the precise mechanism of control, and the extent to which the protein templates are incorporated into the crystals, will require further study (see supplemental discussion).

Our DHR proteins possess greater structural regularity and tunability than previously studied native and engineered biomineralization proteins^14,31,50^ and drive polymorphic specific CaCO_3_ nucleation. This tunability allows rigorous interrogation of the interface driving this templating effect. We found that the spacing of the designed sidechain arrays with similar chemistry had a pronounced effect (Table S2): FD15 with an array spacing of 0.9 nm did not template calcite, DHR49-Neg with a spacing of 1.0 nm templated calcite to some extent, and FD31 with a 1.1 nm spacing was a strong nucleator of calcite. The 1.0 nm and 1.1 nm spacings produced nanocrystals with different preferred orientations, potentially due to the formation of epitaxial matches with different calcite planes (Sup Discussion, Fig. S3, Fig. S14, Fig. S15). We also observed effects of the overall array dimensions: comparing 5, 8 and 11 nm versions of FD31 (FD31-Rep3, FD31, FD31-Rep9, respectively) showed that the smaller templates nucleated larger calcite particles, either because they have higher interfacial free energies and are less effective nucleators, leaving more Ca^2+^ and CO_3_^2-^ available for post-nucleation classical growth, or because they are weaker surface ligands on the nucleated crystals (there is a linear scaling between inverse particle surface area and protein surface area (Fig. S16). Given that template-mineral binding is directly related to the interfacial free energy between the template and the crystal^51^, these two effects are expected to go hand in hand ^52^. The ability to change the surface chemistry simply by recoding the synthetic gene encoding the designs enabled interrogation of the effect of surface chemistry on nucleation outcomes. The observation that FD31-Gln-Checker nucleates calcite better than FD31-Asp suggests that stereochemical alignment is more important to the calcite promoting activity than the total number of negative moieties. Steric hindrance and/or the inclusion of positive moieties within the template may explain the inability of FD31-Lys-Checker to nucleate calcite.

Our results demonstrate that computational protein design now allows the control of proteins over mineral formation that occurs during biomineralization to be studied with genetically encoded molecular templates that have known stable 3D structures, are chemically and structurally tunable, and can be engineered with atom-scale precision ^38,53,54^. Compared to previous studies of natural and engineered proteins^14,15,31^, using de novo designed proteins allows more rigorous testing of how structurally defined biomolecular surfaces control inorganic crystallization through systematic variations in the net charge, pattern of charges, side chain identity, solvent accessible surface area, and surface hydrophobicity. Our approach could lead to routes to enhancing biogenic formation of CaCO3, an important carbon sink in the biogeochemical cycle ^55^. This new approach sets the stage for the programmable control of crystal polymorph, nucleation pathway, growth mechanism, and final crystal morphology, for the ultimate goal of developing hybrid materials with novel functions, including carbon sequestration.

## Acknowledgements

This research was performed at Pacific Northwest National Laboratory (PNNL) with support from the U.S. Department of Energy (DOE), Office of Science (SC), Basic Energy Sciences (BES), Division of Materials Sciences and Engineering, under Award FWP 67554. PNNL is operated by Battelle for the Department of Energy under contract no. DE-AC05-76RLO1830. ATR-FTIR and high-resolution TEM measurements were performed under user proposals 60575 at the Environmental Molecular Sciences Laboratory (EMSL), a DOE SC User Facility sponsored by the Office of Environmental and Biological Research under Contract No. DEAC05-76RL01830. This work is also supported by the US DOE, SC, BES, as part of the Energy Frontier Research Centers program: CSSAS, The Center for the Science of Synthesis Across Scales under Award Number DE-SC0019288.

XRD data were obtained at the Advanced Light Source (ALS) of Lawrence Berkeley National Laboratory (LBNL), which is supported by the U.S. DOE OBES under Contract No. DE-AC02-05CH11231.

The AFM experiments were conducted at the Molecular Analysis Facility, a National Nanotechnology Coordinated Infrastructure (NNCI) site at the University of Washington, which is supported in part by funds from the National Science Foundation (awards NNCI-2025489, NNCI-1542101), the Molecular Engineering & Sciences Institute, and the Clean Energy Institute. The AFM experiments were supported by CSSAS.

This work was supported by DOE Office of Science /DE-FOA-0001664 “Principles of De Novo Protein Nanomaterial Assembly in 1, 2 and 3 Dimensions”; by the grant DE-SC0018940 funded by the U.S. Department of Energy, Office of Science and by the Audacious Project.

