## Supplemental Materials for "Directing polymorph specific calcium carbonate formation with de novo protein templates"

### Materials and Methods

Calcium chloride dihydrate ($CaCl_2$•$2H_2O$, Sigma-Aldrich), sodium bicarbonate ($NaHCO_3$, Sigma-Aldrich), and ultrapure Milli-Q water ($H_2O$, 18.2 MΩ $cm^{-1}$) were used as the reagents for $CaCO_3$ synthesis experiments. Nuclease-free water was bought from Ambion. Tris buffer (pH 7) and Potassium Chloride (KCl) were bought from Sigma-Aldrich. Mica was bought from Ted Pella, CA.

#### Protein design

**FD31**: Protein design was carried out using the Rosetta Macromolecular Modeling Suite(*1*). The protocol used is adapted from Brunette et al.(*2*), and Hicks et al.(*3*), where initial structures were generated using RosettaRemodel followed by scoring with a coarse-grained function weighted with target helical parameters. We tested a number of combinations of sampled fragment lengths of helices and loops to then determine that the best was to sample helices of equal length or with a difference in length within three residues. Helix lengths of 16 to 30 residues were sampled. The helical parameters were biased to have a radius of 500 nm, a rise of 0 Å and a curvature of 0 rad, yielding a protein with a flat, repetitive surface. Inter-repeat distance was biased to 10.9 Å using harmonic constraints and allowing for a deviation of 0.05 Å. Sequence design was carried out using FastDesign with an enforced inner repeat symmetry and enforced interhelical repeat spacing preventing the backbone minimization cycles to create scaffolds with spacings deviating from the target. Residues on one side of the surface of the protein were mutated manually or using a custom PyRosetta protocol to glutamates.

**DHR49-Neg**: The molecule named DHR49 found in the published work of Brunette et al.(*2*) was used as a scaffold. It was turned into DHR49-neg by expanding it to 6 repeat units, and mutating one of its helical surfaces to contain only negatively charged residues (24 aspartates and 18 glutamates). Lastly the capping features were removed by mutating the sequence of the terminal repeats to the sequence of the internal repeats to produce head-to-tail interfaces containing the same hydrophobic interactions found in the core of the protein.

**For FD15**: This molecule is also reported in Huddy et al. (*4*). The design method is reported in the previous citation and is briefly described below: A helical secondary structure element is placed along additional secondary structure elements that will be part of the repeating unit using BundleGridSampler mover in RosettaScripts. The rigid body transformation for the repeat propagation is set by translating a copy of the original helix; copies are added as needed. Degrees of freedom are limited to the helix phase, displacement of repeating helices on the XY plane, and change in height between adjacent helices. In the case of this molecule, the inter-repeat distance was set to 8.7 Å. Sequence design was carried out using FastDesign with an enforced inner repeat symmetry. Residues on one side of the surface of the protein were mutated manually or using a custom PyRosetta protocol to glutamates.

The top protein designs were selected based on various factors such as protein energetics, core packing, secondary structure shape complementarity, helix quality, number of secondary structures in contact, and buried unsatisfied hydrogen bonds. These designs were then subjected to forward folding simulations to identify those that exhibited a funnel shape leading towards a set of low energy relaxed models near the original model, as evidenced by a low root-mean-square deviation (RMSD). Another metric, the area to the left of the folding funnel from the lowest energy point to +8 rosetta energy units, may also be used to select designs with a value less than 25, as previously published(*3*, *5*). Scripts implementing these protocols are available in the GitHub repository associated with this work (https://github.com/fatimadavila/DHR_CaCO3).

### Designed protein flattening protocol

We used a RosettaScripts protocol to create idealized (perfectly flat and repetitive) models of FD15, DHR49-neg, and FD31 with a range of inter-repeat spacings to produce models suited for computationally screening for geometric matches between the proteins and calcite surfaces. First the following constraints are applied: distance constraints to enforce distances between alpha-carbons in adjacent repeats (ranging from 8Å to 15Å with 0.1Å increments), angle constraints to eliminate any curvature (by constraining the angle between repeated alpha-carbons in the first, middle, and last repeat is 180°), and dihedral constraints to prevent any twist (by constraining the torsion angle between two non-repeat c-alphas in the first repeat to and the corresponding two atoms in the last repeat to be 0°). Using a standard Rosetta score function (beta_nov16) with the addition of these constraints to score the models, we performed Monte Carlo sampling of backbone torsion angles while enforcing symmetry among the repeats to find the lowest energy model that satisfies these constraints, and thereby perfectly flat and repetitive. We determined the repeat distance that produced the lowest energy model for each DHR (8.7Å for FD15, 10.6Å for DHR49-neg, and 11.4Å for FD31), and selected the models with repeat distances constrained to within 1Å of these values for docking on calcite surfaces. RosettaScript XML files implementing the protocol  and the resulting models are available on GitHub. (https://github.com/fatimadavila/DHR_CaCO3).

### Docking protocol on calcium carbonate surfaces

The idealized DHR models were then placed onto models of calcite (110), (202), and (104) facets and docked with Rosetta. Each docking trajectory starts by randomly rotating the protein on the surface. This starting pose is input into a Monte Carlo protocol that samples the rigid body orientation of the protein relative to the surface and interfacial side chain rotamers. The side chains are sampled in such a way that the same rotamer is placed at every equivalent repeat position. Finally, the binding score in Rosetta Energy Units (R.E.U.) is calculated by subtracting the median score of models of the protein and surface separated by 10 nm from the score of the protein docked onto the surface. Implementations of this Rosetta protocol and the resulting models are available on GitHub. (https://github.com/fatimadavila/DHR_CaCO3).

### Expression and purification of proteins

Genes encoding the designs were then ordered through Genscript. Constructs with a N-terminal His6-tag followed by a TEV Cleavage site were cloned into either pET-28b+ or pet21b between NdeI and XhoI sites. An additional flexible linker with a tryptophan was added to help with protein concentration determination by absorbance at 280 nm. The cloned genes were transformed into either Lemo21(DE3) E. coli from New England Biolabs (NEB) or in BLR(DE3) E. coli cells from Novagen. Expression then proceeded for 24 hours at 37 ºC using 0.5 L cultures in 2L flasks using Studiers M2 autoinduction media with 50 µg/mL kanamycin or 50 µg/mL carbenicillin for pET-28b+ or pet21b, respectively. Cells were pelleted at 4000 g for 30 minutes at 12 ºC, then resuspended in ~40 mL of lysis buffer (20 mM Tris, 500 mM NaCl, 30 mM Imidazole, 0.25% Chaps, 1mM PMSF,  1 mg/mL DNAse, pH 8) and finally lysed after homogenization using a microfluidizer (Microfluidics M110P) at 18K pounds force per square inch. The lysate was clarified at 24000g for 30 minutes at 12 ºC, and the soluble fraction was filtered through 0.7 µm syringe filters and set to do overnight batch binding with 1.5 mL of Ni-NTA resin (Qiagen) equilibrated in wash buffer (20 mM Tris, 500 mM NaCl, 30 mM Imidazole, 0.25% Chaps, 5% glycerol, pH 8). This was then transferred to a gravity column and washed with 25 mL of wash buffer before elution in 3 mL of elution buffer (20 mM Tris, 500 mM NaCl, 500 mM Imidazole, 0.25% Chaps, 5% glycerol, pH 8). Eluate was then dialysed in 3.5 kDa molecular weight cut-off dialysis cassettes (Thermo) into 5 L of TEV cleavage buffer (50 mM Tris, 50 mM NaCl, pH 8) three times before starting overnight cleavage by adding TEV protease in a ratio of 1 mg for each 25 mg of tagged protein. Secondary IMAC was carried out to remove the TEV protease and uncleaved product. The flowthrough was collected for fractionation by size exclusion chromatography with an AKTA pure chromatography system on a Superdex 200 Increase 10/300 GL column in TBS (20 mM Tris pH 8.0, 100 mM NaCl). The purified proteins were then dialyzed into MOPS buffer (10 mM) adjusted to pH 7. Dialysis was carried out three times overnight with a dialysis ratio of 1:10000 volume each time and aliquots of 20 µl were snap-frozen for long-term storage. DHR49-neg was purified using a different protocol described in Jin et al.(*6*).

### Characterization of Designed Helical Repeat Proteins

#### Measuring protein concentration

Absorbance at 280 nm wavelength of 2 µl of protein samples was measured using a Nanodrop 8000 spectrometer (Thermo Scientific). The concentration was then calculated based on the measured absorbance and the known extinction coefficient following the Beer-Lambert law.

#### Circular Dichroism

Using a Jasco J-1500 CD spectrometer, measurements were taken on a sample with a concentration of 0.3 mg/mL in 20 mM Tris pH 8 and 100 mM NaCl, using a 1 mm path length cuvette. The raw CD signal was divided by N x C x L x 10 to convert it to mean residue ellipticity, where N is the number of residues, C is the protein concentration, and L is the path length of 0.1 cm.

#### Small Angle X-ray Scattering

The SIBYLS High Throughput SAXS Advanced Light Source in Berkeley, California(*7*)was used to collect data. Each sample was exposed to the beam for 0.3 s for a total of 10.2 s, which resulted in 33 frames per sample. The data was collected at both low (~1 mg/mL) and high (~5 mg/mL) protein concentrations in SAXS buffer (25 mM Tris pH 8.0, 150 mM NaCl, 2% glycerol). The SIBYLS website's "SAXS FrameSlice" tool(*8*) was used to analyze the data for the high and low concentration samples and determine the best dataset. If there was evidence of aggregation in any of the 33 frames, only the data points before aggregation occurred were used in the Gunier region; otherwise, all data was included. All data was used for Porod and Wide regions. The resulting dataset was then compared to the predicted SAXS profile generated from the design model using the FoxS SAXS server(*9*).

### Crystallographic analysis

Crystallization experiments were conducted using the sitting drop vapor diffusion method and crystallization trials were set up in 200 nL drops using the 96-well plate format at 20 ˚C. Crystallization plates were set up using a Mosquito LCP from SPT Labtech, then imaged using UVEX microscopes and UVEX PS-256 from JAN Scientific. Diffraction quality crystals formed in 0.1 M TRIS pH 6.5, and 25% w/v Polyethylene glycol 3,350. Crystals were flash frozen in liquid nitrogen before sending them to the synchrotron.
Diffraction data was collected at the Advanced Photon Source beamline on 24-ID-C. X-ray intensities and data reduction were evaluated and integrated using XDS(*10*) and merged/scaled using Pointless/Aimless in the CCP4 program suite(*11*). Structure determination and refinement starting phases were obtained by molecular replacement using Phaser(*12*) using the designed model for the structures. Following molecular replacement, the models were improved using phenix.autobuild(*13*); efforts were made to reduce model bias by setting rebuild-in-place to false, and using simulated annealing and prime-and-switch phasing. Structures were refined in Phenix(*13*). Model building was performed using COOT(*14*).

### $CaCO_3$ crystallization experiments

In a typical crystallization experiment, 0.5 ml 10 mM $CaCl_2$ was mixed with 10 mM $NaHCO_3$ as a control group. In the experimental group, we mixed 10 mM $CaCl_2$ with 2.16 μM first and then added 10 mM $NaHCO_3$ to initiate the nucleation and growth of $CaCO_3$.

### Ex situ TEM

Ex-situ TEM samples were prepared by dropping 0.6 μl reaction solutions collected over a range of time points on a carbon-coated Cu-grid (300 mesh, purchased from Ted Pella) which was treated by plasma cleaning. TEM was performed in a FEI Titan ETEM 80–300 kV operated at 300 kV.

### Cryo-TEM

Cryo-TEM experiments were performed in a Titan ETEM 80–300 kV. Prior to the vitrification procedure, a pure lacey carbon grid is surface plasma-treated to make it hydrophilic. Using an automated vitrification robot (FEI Vitrobot Mark III, blot time: 3 s), a 3.0 μl mixed aqueous solution including 5 mM $CaCl_2$ and proteins as well as 5 mM $NaHCO_3$ was loaded onto a grid and plunged into liquid ethane. The frozen sample was saved and transferred in a cryogenic holder and cryo-TEM (FEI ETEM operated at 300 kV) under liquid nitrogen conditions, and thus the microscope maintained its temperature near −192 °C throughout the holder.

### UV-Vis and DLS

The $CaCl_2$ incubated FD31 protein solution was measured by UV-Vis spectrophotometer (Ultrospec 2100 pro) and dynamic light scattering (DLS, using a Malvern Zetasizer) to evaluate the protein-Ca interactions.

### In situ ATR-FTIR

The crystallization process of $CaCO_3$ was monitored in situ using a Bruker LUMOS II Fourier transform infrared (FTIR) spectrometer with Superior μ-attenuated total reflectance (ATR) FT-IR Capabilities. The retractable diamond crystal is controlled by high-precision piezoelectric motors and integrated into the lens, which allows us to precisely control the detector location in the reaction solution. The crystallization experiments were initiated by adding 200 μl 10 mM $NaHCO_3$ into 200 μl 10 mM $CaCl_2$ with or without DHR proteins. For each FTIR spectrum recorded, 8 scans were carried out at a resolution of 2 $cm^{-1}$ using $H_2O$ as background. The first spectrum was recorded after mixing the solutions for ~5 s.

### LP-TEM

All of the liquid-cell chips (Hummingbird Scientific), consisted of two square 4 $mm^2$ silicon chips with 50 nm thick silicon nitride ($Si_3N_4$) membranes in 50 × 200 $\mu m^2$ windows for imaging, were plasma cleaned (Harrick Plasma Cleaner) for 2 min before use. In a typical experiment, 0.3 μl 10 mM $CaCl_2$ containing 2.16 μM FD31 was dropped onto the bottom chip. This was followed by adding 0.3 μl 10 mM $NaHCO_3$ solution, and, finally, the reaction solution was sealed using a window chip. The sealed chips were assembled inside the liquid cell holder (Hummingbird Scientific) and were leak-checked. After that, the holder was immediately inserted into the TEM for observation within ~5 min. TEM was carried out in a field emission Titan ETEM 80–300 kV (Thermo Fisher Scientific) operated at 300 kV. TEM images were acquired using an Eagle CCD (1,024 × 1,024 pixels). In situ movies were recorded using free software called Camstudio for Screen and Video Recorder. All images from in situ movies were processed using the open-source software ImageJ. To minimize beam effects, a low electron dose rate (~100 $e/nm^2{\bullet}s$) was used to observe the formation process by adjusting the condenser aperture size (50 μm), and spot size (3).

### AFM

For DHR49-Neg, the protein stock solution was diluted to 0.5 μM in a 20 mM Tris buffer with 3M KCl. Then 100 μl diluted protein solution was incubated on freshly cleaved mica for 30 min. For FD31, the protein stock solution was diluted to 1.0 μM in nuclease-free water with 5 mM $CaCl_2$. Then 100 μl diluted protein solution was incubated on freshly cleaved mica for 10 min. The as-assembled proteins on mica were imaged using Cypher-ES AFM (Asylum Research, CA) in a 20 mM Tris buffer with 3 M KCl, and 5 mM $CaCl_2$, respectively. The amplitude modulation mode and SNL-10-A probe (Bruker, CA) were used in the AFM experiments. The data processing was done with SPIP software (Image Metrology, Denmark).

### Supplementary text

#### Number density estimation

Given calcite density: 2.7 g/cm$^3$, Mw=100 g/mol, and assuming all precursor $CaCO_3$ (1 ml 5 mM) is transformed into calcite because of its very low solubility (~1.3 μM), it will produce CaCO3 solids with a total volume of 1.85E17 nm$^3$. The average cubic-like calcite nanoparticles length= 5.5 nm (Figure 5b), which gives a volume of ~166 nm$^3$, enabling us estimate the number of calcite nanocrystals to be 1.1E15, which is comparable to the number of protein monomers: 6.5E14 given 1 ml 1.08 uM FD31, and $N_A$=6E23.

#### The electron beam or confinement effects in LP-TEM experiments

To confirm that the observed nucleation of calcite nanocrystals did not result from the continuous exposure to the electron beam or confinement effects, but solely from the interplay between the $CO_3^{2-}$ and the protein-Ca complex, the reaction solution was imaged in a neighboring area without an electron beam irradiation. Calcite nanocrystals with the average size of ~5.5 nm were identified (Fig. 5b). These newly formed calcite nanocrystals keep moving in solution and tend to aggregate, rather than dissolve (Fig. S10). It is plausible that the negatively charged -$COO^-$ in the proteins are absorbed on the calcite nanocrystals, as there are observations of low contrast material on the surface of the nanocrystals (Fig. S3c), thus stabilizing them. This is supported by the fact that we failed to observe the gradual growth into bigger rhombohedral calcite crystals as seen in previous in situ TEM observations(*15*). Further benchtop experiments providing TEM, Cryo-TEM, and in situ liquid-phase ATR-FTIR data confirmed these results for equivalent bulk solutions. We do not expect confinement effects in LP-TEM experiments to affect nucleation rates or pathways since the dimensions for both the calcite nanocrystals (considering a critical nucleus size of ~1-5 nm) and the protein-Ca complex are small compared to those of the liquid-cell.

#### Proposed mechanisms of protein control over calcite nucleation

Although the precise mechanism of control over nucleation cannot be discerned from the data presented here, given the direct appearance of calcite for certain distributions of carboxylic side chains, the implication is that those distributions bias the configuration of the ions at the protein-solution interface towards that of the calcite lattice. Moreover, because the free energy barrier and critical nucleus size are determined by the probability that ions achieve a size and configuration for which subsequent growth decreases the free energy, this implication is consistent with the rationale originally used for selecting the DHR protein design with its large flat surface and periodic distribution of carboxylic groups.

A variety of factors may explain the different effects of the different proteins. Firstly, the use of odd (Asp) vs. even (Glu) number of carbons in the side chains of the binding moieties could bias the formation of different interfaces by virtue of the angle at which the carboxyl group is displayed on the surface of the protein, as seen self-assembled monolayers(*16*). If the preferred angle of the carboxyl group stereochemically matches the orientation of the carbonate groups on that facet, the interaction will be favored. Glu side chains better match (110) and Asp better matches (202)(Fig. S9). Observations that FD31, which exclusively presents Glu, stabilizes the (110) orientation of calcite nanoparticles, whereas DHR49-Neg includes Asp residues and yields (202) orientations, suggesting a stereochemical explanation for these differences.

Secondly, geometric lattice matching may play a role. DHRs with different repeat spacings (Fig. S4a-c) affect $CaCO_3$ growth differently (Fig. 2), and suggest interactions between specific proteins and specific interfaces (Fig. 4, Fig. S8). Rosetta docking simulations were run with DHR models constrained to be flat and repetitive to identify potential geometric matches at the protein-mineral interfaces (see methods). Since the constrained DHR models with repeat spacings within a ~0.5 Å of the minima had comparable predicted energies (Fig. S4d), models of FD15, DHR49-neg, and FD31 with repeat distances constrained to a range of values surrounding the minima (to represent the possibility of small conformational changes in the proteins) were docked onto models of three calcite facets while maintaining repeat symmetry. The (110) and (202) calcite facets were included based on the observed orientations of nanoparticles formed in the presence of FD31 and DHR49-Neg, respectively, and the (104) surface was included as a control because it is typically expressed in calcite crystals. This Rosetta modeling does not accurately represent the complex energetics of protein-mineral interactions at solid-liquid interfaces, discriminate between proteins that nucleate or do not nucleate, or explain facet specificity. However, it was consistent with observations that these DHR proteins bind (110) and (202) rather than the (104) facet (Fig. S15) and yielded lattice-matching docked conformations of the proteins on the surfaces (Fig. S16). At minimum these models illustrate how DHR proteins are structurally well suited to epitaxially template the growth $CaCO_3$, and to our knowledge they are among the best supported models of protein-mineral interfaces that drive heterogeneous nucleation currently available. These models are provided in a GitHub repository alongside the scripts that produced them (https://github.com/fatimadavila/DHR_CaCO3).

### Incorporation of proteins into calcite crystals

The results also leave undetermined the extent to which the protein templates are incorporated into the crystals or driven to detach from the nano-calcite surfaces as the crystals assemble. The observation of continuous lattice fringes between attached particles in a number of instances (Figures 4c and S7h), as well as a significant number of nanocrystals that continue to attach to mature micron-scale calcite crystals (Figure S10e), suggests that at least some proteins do detach and can again act as a template to generate new nano-calcite crystals. In that regard, the protein templates may be thought of as analogous to heterogeneous catalysts:

they act as surfaces that reduce the energy barrier to form calcite from $Ca^{2+}$and $CO_3^{2-}$ ions, but are not consumed in the process.

### Supporting tables

**Table S1: Sequences of tested proteins. *Indicates sequences with high expression levels. **Indicates sequences that were further characterized and tested in nucleation trials. ***Indicates variants of FD31 that were tested for nucleation activity. Underlined sequences were removed during purification by cleavage with TEV protease.**

| Name | Sequence |
|---|---|
| FD15** | MGSSHHHHHHSSGLVPRGSHMENLYFQGSWSGGSGGEAADEARRAIEAALEEARAAADEARSDSTGETVKKAVDKAEKAAEDAFREIKQAVNQAEKQGASEAAFEAFAAIAAAAAEAAAAAFEAFSDSTGETVAEAVAKALKAAMEAFAEIAKAVAQAAKQGASEAAFEAFAAIAAAAAEAAAAAFEAFSDSTGETVAEAVAKALKAAMEAFAEIAKAVAQAAKQGASEAAFEAFAAIAAAAAEAAAAAFEAFSDSTGETVAEAVAKALKAAMEAFAEIAKAVAQAAKQGASEAAFEAFAAIAAAAAEAAAAAFEAFSDSTGETVAEAVAKALKAAMEAFAEIAKAVAQAAKQGASEEAFEKFAAIAAEAAEAAAAAFERFSDSTGETEAEKVAKELKQLMEEFAERAKSVAEQAKNGAS |
| DHR49-Neg** | MDSKVLEEAIRVIAEIAKESGSEDAAESAIDAVADIADEAQDSKVLEEAIRVIAEIAKESGSEDAAESAIDAVADIADEAQDSKVLEEAIRVIAEIAKESGSEDAAESAIDAVADIADEAQDSKVLEEAIRVIAEIAKESGSEDAAESAIDAVADIADEAQDSKVLEEAIRVIAEIAKESGSEDAAESAIDAVADIADEAQDSKVLEEAIRVIAEIAKESGSEDAAESAIDAVADIADEAQGW |
| FD31** | MGSSHHHHHHSSGLVPRGSHMENLYFQGSWSGGSGGPEEALEEVEERIEELESALESNPTNEEELREILKKILEIFEELFREAKARNDTELLLEAVEAVIELLETLLELNPTNEELLREILKIILRIFELLFELAKKQNDTELLLEAVEAVIELLETLLELNPTNEELLREILKIILRIFELLFELAKKQNDTELLLEAVEAVIELLETLLELNPTNEELLREILKIILRIFELLFELAKKQNDTELLLEAVEAVIELLETLLELNPTNEELLREILKIILRIFELLFELAKKQNDTELLSEAKEAVAELLETLAELNPTNQELKEEIKKIQERIAELEKELAEKQNA |
| FD31-3rep*** | MGSSHHHHHHSSGLVPRGSHMENLYFQGSWSGGSGGPEEALEEVEERIEELESALESNPTNEEELREILKKILEIFEELFREAKARNDTELLLEAVEAVIELLETLLELNPTNEELLREILKIILRIFELLFELAKKQNDTELLSEAKEAVAELLETLAELNPTNQELKEEIKKIQERIAELEKELAEKQNA |
| FD31-9rep*** | MGSSHHHHHHSSGLVPRGSHMENLYFQGSWSGGSGGPEEASEEVEERSEELESALESNPTNEEELREILKKIQEIFEELFREAKARNDTELLSEAVEAVSELLETLLENNPTNEELLREILKIIQRIFELLFELAKKQNDTELLSEAVEAVSELLETLLENNPTNEELLREILKIIQRIFELLFELAKKQNDTELLSEAVEAVSELLETLLENNPTNEELLREILKIIQRIFELLFELAKKQNDTELLSEAVEAVSELLETLLENNPTNEELLREILKIIQRIFELLFELAKKQNDTELLSEAVEAVSELLETLLENNPTNEELLREILKIIQRIFELLFELAKKQNDTELLSEAVEAVSELLETLLENNPTNEELLREILKIIQRIFELLFELAKKQNDTELLSEAKEAVSELLETLAENNPTNQELKEEIKKIQERIAELEKELAEKQNA |
| FD31-Asp*** | MGSSHHHHHHSSGLVPRGSHMENLYFQGSWSGGSGGPEEALDEVDERIDELESALDSNPTNEEELREILKKILEIFEELFREAKARNDTELLLDAVDAVIDLLETLLDLNPTNEELLREILKIILRIFELLFELAKKQNDTELLLDAVDAVIDLLETLLDLNPTNEELLREILKIILRIFELLFELAKKQNDTELLLDAVDAAIDLLETLLDLNPTNEELLREILKIILRIFELLFELAKKQNDTELLSDAKDAVADLLETLADLNPTNQELKEEIKKIQERIAELEKELAEKQNA |

| | |
|---|---|
| FD31-Gln-Checker*** | MGSSHHHHHHSSGLVPRGSHMENLYFQGSWSGGSGGPEEALQEVEERIQELESALQSNPTNEEELREILKKILEIFEELFREAKARNDTQLLLEAVQAVIELLQTLLELNPTNEELLREILKIILRIFELLFELAKKQNDTELLLQAVEAVIQLLETLLQLNPTNEELLREILKIILRIFELLFELAKKQNDTQLLLEAVQAVIELLQTLLELNPTNEELLREILKIILRIFELLFELAKKQNDTELLLQAVEAAIQLLETLLQLNPTNEELLREILKIILRIFELLFELAKKQNDTQLLSEAKQAVAELLQTLAELNPTNQELKEEIKKIQERIAELEKELAEKQNA |
| FD31-Lys-Checker*** | MGSSHHHHHHSSGLVPRGSHMENLYFQGSWSGGSGGPEEALKEVEERIEELKSALESNPTNEEELREILKKILEIFEELFREAKARNDTKLLLEAVKAVIELLETLLKLNPTNEELLREILKIILRIFELLFELAKKQNDTELLLEAVEAVIKLLETLLELNPTNEELLREILKIILRIFELLFELAKKQNDTELLLKAVEAVIELLKTLLELNPTNEELLREILKIILRIFELLFELAKKQNDTKLLLEAVKAAIELLETLLKLNPTNEELLREILKIILRIFELLFELAKKQNDTELLSEAKEAVAKLLETLAELNPTNQELKEEIKKIQERIAELEKELAEKQNA |
| FD01 | MGSSHHHHHHSSGLVPRGSHMENLYFQGSWSGGSGGPEELLREVRERIEELKSLLESNPTNEELLREILKKILEIFEELFREAKARNDVELLLRAVRLVIELLKTLLELNPTNEELLREILKIILRIFELLFELAKKLNDVELLLRAVRLVIELLKTLLELNPTNEELLREILKIILRIFELLFELAKKLNDVELLLRAVRLVIELLKTLLELNPTNEELLREILKIILRIFELLFELAKKLNDVELLLRAVRLVIELLKTLLELNPTNEELLREILKIILRIFELLFELAKKLNDTELLSRAKELVAELLRTLLELNPTNQELLEEIKKIQERIAELEKELLEKLNA |
| FD02 | MGSSHHHHHHSSGLVPRGSHMENLYFQGSWSGGSGGPEELLAEVESRIAELESLLASNPTNEELLREILKKILEIFEELFREAKARNDVELLLAAVELVIALLETLLALNPTNEELLREILKIILRIFELLFELAKKLNDVELLLAAVELVIALLETLLALNPTNEELLREILKIILRIFELLFELAKKLNDVELLLAAVELVIALLETLLALNPTNEELLREILKIILRIFELLFELAKKLNDVELLLAAVELVIALLETLLALNPTNEELLREILKIILRIFELLFELAKKLNDTELLSAAKELVAELLETLLELNPTNQELLEEIKKIQERIAELEKELLEKLNA |
| FD03* | MGSSHHHHHHSSGLVPRGSHMENLYFQGSWSGGSGGPEELLREILKIILRIFELLFELAKKLNDVELLLAAVELVIALLETLLALNPTNEELLREILKIILRIFELLFELAKKLNDVELLLAAVELVIALLETLLALNPTNEELLREILKIILRIFELLFELAKKLNDVELLLAAVELVIALLETLLALNPTNEELLREILKIILRIFELLFELAKKLNDVELLLAAVELVIALLETLLALNP |
| FD04 | MGSSHHHHHHSSGLVPRGSHMENLYFQGSWSGGSGGPELLLAAVEAVIAALETLLALNPTNEELLREILKIILRIFEALFELAKKLNDVELLLAAVELVIALLETLLALNPTNEELLREILKIILRIFELLFELAKKLNDVELLLAAVELVIALLETLLALNPTNEELLREILKIILRIFELLFELAKKLNDVELLAAAVELAAALLETLLALNPTNEELLREILKIIARIAELLAELAKKLN |
| FD05 | MGSSHHHHHHSSGLVPRGSHMENLYFQGSWSGGSGGPAELLEEVESRIEELESLLASNPTNEELLREILKKILEIFEELFREAKARNDVALLLEAVELVIELLETLLALNPTNEELLREILKIILRIFELLFELAKKLNDVALLLEAVELVIELLETLLALNPTNEELLREILKIILRIFELLFELAKKLNDVALLLEAVELVIELLETLLALNPTNEELLREILKIILRIFELLFELAKKLNDTALLSEAIELVAELLETLLALNPTNQELLEEIKKIQERIAELEKELLEKLNA |
| FD06 | MGSSHHHHHHSSGLVPRGSHMENLYFQGSWSGGSGGPEELLREILKIILRIFELLFELAKKLNDVALLLEAVELVIELLETLLALNPTNEELLREILKIILRIFELLFELAKKLNDVALLLEAVELVIELLETLLALNPTNEELLREILKIILRIFELLFELAKKLNDVALLLEAVELVIELLETLLALNP |

| | |
|---|---|
| FD07 | MGSSHHHHHHSSGLVPRGSHMENLYFQGSWSGGSGGPALLLEAVEAVIEALETLLALNPTNEELLREILKIILRIFEALFELAKKLNDVALLLEAVELVIELLETLLALNPTNEELLREILKIILRIFELLFELAKKLNDVALLLEAVELVIELLETLLALNPTNEELLREILKIILRIFELLFELAKKLNDVALLAEAVELAAELLETLLALNPTNEELLREILKIIARIAELLAELAKKLN |
| FD08* | MGSSHHHHHHSSGLVPRGSHMENLYFQGSWSGGSGGPALLLEAVEAVIEALETLLALNPTNEELLREILKIILRIFEALFELAKKLNDVALLLEAVELVIELLETLLALNPTNEELLREILKIILRIFELLFELAKKLNDVALLLEAVELVIELLETLLALNPTNEELLREILKIILRIFELLFELAKKLNDVALLAEAVELAIELLETLLALNPTNEELLREIAKIIARIFELLAELAKKLN |
| FD09 | MGSSHHHHHHSSGLVPRGSHMENLYFQGSWSGGSGGPEELLEEVESRIEELESLLESNPTNEELLREILKKILEIFEELFREAKARNDVELLLEAVELVIELLETLLELNPTNEELLREILKIILRIFELLFELAKKLNDVELLLEAVELVIELLETLLELNPTNEELLREILKIILRIFELLFELAKKLNDVELLLEAVELVIELLETLLELNPTNEELLREILKIILRIFELLFELAKKLNDVELLLEAVELVIELLETLLELNPTNEELLREILKIILRIFELLFELAKKLNDTELLSEAIELVAELLETLLELNPTNQELLEEIKKIQERIAELEKELLEKLNA |
| FD10 | MGSSHHHHHHSSGLVPRGSHMENLYFQGSWSGGSGGPEELLAEVESRIAELESLLESNPTNEELLREILKKILEIFEELFREAKARNDVELLLAAVELVIALLETLLELNPTNEELLREILKIILRIFELLFELAKKLNDVELLLAAVELVIALLETLLELNPTNEELLREILKIILRIFELLFELAKKLNDVELLLAAVELVIALLETLLELNPTNEELLREILKIILRIFELLFELAKKLNDVELLLAAVELVIALLETLLELNPTNEELLREILKIILRIFELLFELAKKLNDTELLSAAIELVAALLETLLELNPTNQELLEEIKKIQERIAELEKELLEKLNA |
| FD11 | MGSSHHHHHHSSGLVPRGSHMENLYFQGSWSGGSGGPDELLDEVDSRIDELDSLLDSNPTNEELLREILKKILEIFEELFREAKARNDVDLLLDAVDLVIDLLDTLLDLNPTNEELLREILKIILRIFELLFELAKKLNDVDLLLDAVDLVIDLLDTLLDLNPTNEELLREILKIILRIFELLFELAKKLNDVDLLLDAVDLVIDLLDTLLDLNPTNEELLREILKIILRIFELLFELAKKLNDVDLLLDAVDLVIDLLDTLLDLNPTNEELLREILKIILRIFELLFELAKKLNDTDLLSDAIDLVADLLDTLLDLNPTNQELLEEIKKIQERIAELEKELLEKLNA |
| FD12 | MGSSHHHHHHSSGLVPRGSHMENLYFQGSWSGGSGGPDELLAEVDSRIAELDSLLDSNPTNEELLREILKKILEIFEELFREAKARNDVDLLLAAVDLVIALLDTLLDLNPTNEELLREILKIILRIFELLFELAKKLNDVDLLLAAVDLVIALLDTLLDLNPTNEELLREILKIILRIFELLFELAKKLNDVDLLLAAVDLVIALLDTLLDLNPTNEELLREILKIILRIFELLFELAKKLNDVDLLLAAVDLVIALLDTLLDLNPTNEELLREILKIILRIFELLFELAKKLNDTDLLSAAIDLVAALLDTLLDLNPTNQELLEEIKKIQERIAELEKELLEKLNA |
| FD13 | MGSSHHHHHHSSGLVPRGSHMENLYFQGSWSGGSGGPAELLDEVDSRIDELDSLLASNPTNEELLREILKKILEIFEELFREAKARNDVALLLDAVDLVIDLLDTLLALNPTNEELLREILKIILRIFELLFELAKKLNDVALLLDAVDLVIDLLDTLLALNPTNEELLREILKIILRIFELLFELAKKLNDVALLLDAVDLVIDLLDTLLALNPTNEELLREILKIILRIFELLFELAKKLNDVALLLDAVDLVIDLLDTLLALNPTNEELLREILKIILRIFELLFELAKKLNDTALLSDAIDLVADLLDTLLALNPTNQELLEEIKKIQERIAELEKELLEKLNA |
| FD14 | MGSSHHHHHHSSGLVPRGSHMENLYFQGSWSGGSGGPEELLREVRLRIRELKSLLRSNPTNEELLREILKKILEIFEELFREAKARNDVELLLRAVRLVIRLLKTLLRLNPTNEELLREILKIILRIFELLFELAKKLNDVELLLRAVRLVIRLLKTLLRLNPTNEELLREILKIILRIFELLFELAKKLNDVELLLRAVRLVIRLLKTLLRLNPTNEELLREILKIILRIFELLFELAKKLNDVELLLRAVRLVIRLLKTLLRLNPTNEELLREILKIILRIFELLFELAKKLNDTELLSRAKRLVARLLKTLLRLNPTNQELLEEIKKIQERIAELEKELLEKLNA |

| | |
|---|---|
| FD16* | MGSSHHHHHHSSGLVPRGSHMENLYFQGSWSGGSGGDAEEAAREAEAAIKEAEDEAREAGASEEALKAAKQAFDEIRKAMREAEKSGASEDALEAAAEAFVAIAEAAAEALEAGASEEALKAAAQAFAEIAKAMAEALKSGASEDALEAAAEAFVAIAEAAAEALEAGASEEALKAAAQAFAEIAKAMAEALKSGASEDALEAAAEAFVAIAEAAAEALEAGASEEALKAAAQAFAEIAKAMAEALKSGASEDALEAAAEAFVAIAEAAAEALEAGASEEALKAAAQAFAEIAKAMAEALKSGASEDALEAAAEEFVRIAEAAAEALEAGASEEELREAAKRFAEEAKRMAEELKSGRSA |
| FD17 | MGSSHHHHHHSSGLVPRGSHMENLYFQGSWSGGSGGDADAELREAEEEAKEAAEEAKSDGGDAEAAKKALEFAKKAFDQLREAAKKGADPDAIAELAEAIAEALEAAAEAFSDGGDAEAAAKALEFAAKAFAQLAEAASKGADPDAIAELAEAIAEALEAAAEAFSDGGDAEAAAKALEFAAKAFAQLAEAASKGADPDAIAELAEAIAEALEAAAEAFSDGGDAEAAAKALEFAAKAFAQLAEAASKGADPDAIAELAEAIAEALEAAAEAFSDGGDAEAAAKALEFAAKAFAQLAEAASKGADPDAIASLAEAIAEALERAAETFSDGGDAEEAAKKLEEDAKRFAQKAEEAAKGADP |
| FD18 | MGSSHHHHHHSSGLVPRGSHMENLYFQGSWSGGSGGPEKLASDLEQLAERFEEALARDDEESLRQLLQEFLKIVEKAAKDNNEELLALALELLAELFELALAYNKEELLKLLLQIFLRIVEQAAKAGNEELLALALELLAELFELALAYNKEELLKLLLQIFLRIVEQAAKAGNEELLALALELLAELFELALAYNKEELLKLLLQIFLRIVEQAAKAGNEELLALALELLAELFELALAYNKEELLKLLLQIFLRIVEQAAKAGNEELLALARELLAELQELARAYNKEELLKLLDQIRKRIEDQKRKDGR |
| FD19 | MGSSHHHHHHSSGLVPRGSHMENLYFQGSWSGGSGGQEDLAEDLASTVARAESALEANNNEELRHLSEKLKRLLEELARRGDQELLAELLALIVALAEAALEANNNELLRTLSELLKKLLELLARRGDQELLAELLALIVALAEAALEANNNELLRTLSELLKKLLELLARRGDQELLAELLALIVALAEAALEANNNELLRTLSELLKKLLELLARRGDQELLAELLALIVALAEAALEANNNELLRTLSELLKKLLELLARRGDRELLAELLALISALAEAAQEANNNELKRTREELEKKLSELQSRRGD |
| FD20 | MGSSHHHHHHSSGLVPRGSHMENLYFQGSWSGGSGGREELRQEALRALKDFLKKLEELLARGNREKFAELLASFLEQLEELLAKAFEAGDRELLRQLALEALKIFLKLLELLLARGNRELFAELLALFLALLELLLALAFEAGDRELLRQLALEALKIFLKLLELLLARGNRELFAELLALFLALLELLLALAFEAGDRELLRQLALEALKIFLKLLELLLARGNRELFAELLALFLALLELLLALAFEAGDRELLRQLALEALKIFLKLLELLLARGNRELFAELLALFLALLELLLALAFEAGDRELLRQLAQEARKIFKKLQEELRARGNEELFAELEALFSALEELLDALLQEAGD |
| FD21 | MGSSHHHHHHSSGLVPRGSHMENLYFQGSWSGGSGGREELRQEALRALKDFLKKLEELLARGNREKFAELLASFLEQLEELLAKAFEAGDRELLRQLALEALKIFLKLLELLLARGNRELFAELLALFLELLEELLALAFEAGDRELLRQLALEALKIFLKLLELLLARGNRELFAELLALFLELLEELLALAFEAGDRELLRQLALEALKIFLKLLELLLARGNRELFAELLALFLELLEELLALAFEAGDRELLRQLALEALKIFLKLLELLLARGNRELFAELLALFLELLEELLALAFEAGDRELLRQLAQEARKIFKKLQEELRARGNEELFAELEALFSELEEELDALLQEAGD |
| FD22 | MGSSHHHHHHSSGLVPRGSHMENLYFQGSWSGGSGGREELRQEALRALKDFLKKLEELLARGNREKFAELLASFLEQLEELLAKAFEAGDRELLRQLALEALKIFLKLLELLLARGNRELFAELLALFLALLELLLALAFEAGDRELLRQLALEALKIFLKLLELLLARGNRELFAELLALFLALLELLLALAFEAGDRELLRQLALEALKIFLKLLELLLARGNRELFAELLALFLALLELLLALAFEAGDRELLRQLALEALKIFLKLLELLLARGNRELFAELLALFLALLELLLALAFEAGDRELLRQLAQEARKIFKKLQEELRARGNEELFAELEALFSALEELLDALLQEAGD |

| | |
|---|---|
| FD23* | MGSSHHHHHHSSGLVPRGSHMENLYFQGSWSGGSGGPEEALREVRERIEELKSALESN<br>PTNEEELREILKKILEIFEELFREAKARNDTELLLRAVRAVIELLKTLLELNPTNEELLREILKII<br>LRIFELLFELAKKQNDTELLLRAVRAVIELLKTLLELNPTNEELLREILKIILRIFELLFELAKKQ<br>NDTELLLRAVRAVIELLKTLLELNPTNEELLREILKIILRIFELLFELAKKQNDTELLLRAVRAVI<br>ELLKTLLELNPTNEELLREILKIILRIFELLFELAKKQNDTELLSRAKEAVAELLRTLAELNPTN<br>QELKEEIKKIQERIAELEKELAEKQNA |
| FD24 | MGSSHHHHHHSSGLVPRGSHMENLYFQGSWSGGSGGPEEALREVRERIEELKSALES<br>NPTNEEELREILKKILEIFEELFREAKARNDVELLLRAVRAVIELLKTLLELNPTNEELLREIL<br>KIILRIFELLFELAKKQNDVELLLRAVRAVIELLKTLLELNPTNEELLREILKIILRIFELLFELAK<br>KQNDVELLLRAVRAVIELLKTLLELNPTNEELLREILKIILRIFELLFELAKKQNDVELLLRAV<br>RAVIELLKTLLELNPTNEELLREILKIILRIFELLFELAKKQNDTELLSRAKEAVAELLRTLAEL<br>NPTNQELKEEIKKIQERIAELEKELAEKQNA |
| FD25 | MGSSHHHHHHSSGLVPRGSHMENLYFQGSWSGGSGGPEEALREVRERIEELKSALESN<br>PTNEEELREILKKILEIFEELFREAKARNDTELLLRAVRAVIELLKTLLELNPTNEELLREILKII<br>LRIFELLFELAKKLNDTELLLRAVRAVIELLKTLLELNPTNEELLREILKIILRIFELLFELAKKLN<br>DTELLLRAVRAVIELLKTLLELNPTNEELLREILKIILRIFELLFELAKKLNDTELLLRAVRAVIEL<br>LKTLLELNPTNEELLREILKIILRIFELLFELAKKLNDTELLSRAKEAVAELLRTLAELNPTNQE<br>LKEEIKKIQERIAELEKELAEKLNA |
| FD26 | MGSSHHHHHHSSGLVPRGSHMENLYFQGSWSGGSGGPEEALREVRERIEELKSALESN<br>PTNEEELREILKKIQEIFEELFREAKARNDTELLLRAVRAVIELLKTLLELNPTNEELLREILKII<br>QRIFELLFELAKKQNDTELLLRAVRAVIELLKTLLELNPTNEELLREILKIIQRIFELLFELAKKQ<br>NDTELLLRAVRAVIELLKTLLELNPTNEELLREILKIIQRIFELLFELAKKQNDTELLLRAVRAVI<br>ELLKTLLELNPTNEELLREILKIIQRIFELLFELAKKQNDTELLSRAKEAVAELLRTLAELNPTN<br>QELKEEIKKIQERIAELEKELAEKQNA |
| FD27* | MGSSHHHHHHSSGLVPRGSHMENLYFQGSWSGGSGGPEEALAEVEERIAELESALESN<br>PTNEEELREILKKILEIFEELFREAKARNDTELLLAAVEAVIALLETLLELNPTNEELLREILKIIL<br>RIFELLFELAKKQNDTELLLAAVEAVIALLETLLELNPTNEELLREILKIILRIFELLFELAKKQN<br>DTELLLAAVEAVIALLETLLELNPTNEELLREILKIILRIFELLFELAKKQNDTELLLAAVEAVIAL<br>LETLLELNPTNEELLREILKIILRIFELLFELAKKQNDTELLSAAKEAVAELLETLAELNPTNQE<br>LKEEIKKIQERIAELEKELAEKQNA |
| FD28* | MGSSHHHHHHSSGLVPRGSHMENLYFQGSWSGGSGGPEEALAEVEERIAELESALESN<br>PTNEEELREILKKILEIFEELFREAKARNDVELLLAAVEAVIALLETLLELNPTNEELLREILKII<br>LRIFELLFELAKKQNDVELLLAAVEAVIALLETLLELNPTNEELLREILKIILRIFELLFELAKKQ<br>NDVELLLAAVEAVIALLETLLELNPTNEELLREILKIILRIFELLFELAKKQNDVELLLAAVEAVI<br>ALLETLLELNPTNEELLREILKIILRIFELLFELAKKQNDTELLSAAKEAVAELLETLAELNPTN<br>QELKEEIKKIQERIAELEKELAEKQNA |
| FD29 | MGSSHHHHHHSSGLVPRGSHMENLYFQGSWSGGSGGPEEALAEVEERIAELESALESN<br>PTNEEELREILKKILEIFEELFREAKARNDTELLLAAVEAVIALLETLLELNPTNEELLREILKIIL<br>RIFELLFELAKKLNDTELLLAAVEAVIALLETLLELNPTNEELLREILKIILRIFELLFELAKKLND<br>TELLLAAVEAVIALLETLLELNPTNEELLREILKIILRIFELLFELAKKLNDTELLLAAVEAVIALL<br>ETLLELNPTNEELLREILKIILRIFELLFELAKKLNDTELLSAAKEAVAELLETLAELNPTNQEL<br>KEEIKKIQERIAELEKELAEKLNA |

| | |
|---|---|
| FD30 | MGSSHHHHHHSSGLVPRGSHMENLYFQGSWSGGSGGPEEALAEVEERIAELESALASN<br>PTNEEELREILKKILEIFEELFREAKARNDVELLLAAVEAVIALLETLLALNPTNEELLREILKII<br>LRIFELLFELAKKQNDVELLLAAVEAVIALLETLLALNPTNEELLREILKIILRIFELLFELAKKQ<br>NDVELLLAAVEAVIALLETLLALNPTNEELLREILKIILRIFELLFELAKKQNDVELLLAAVEAVI<br>ALLETLLALNPTNEELLREILKIILRIFELLFELAKKQNDTELLSAAKEAVAELLETLAALNPTN<br>QELKEEIKKIQERIAELEKELAEKQNA |
| FD32 | MGSSHHHHHHSSGLVPRGSHMENLYFQGSWSGGSGGPEEALEEVEERIEELESALESN<br>PTNEEELREILKKILEIFEELFREAKARNDVELLLEAVEAVIELLETLLELNPTNEELLREILKII<br>LRIFELLFELAKKQNDVELLLEAVEAVIELLETLLELNPTNEELLREILKIILRIFELLFELAKKQ<br>NDVELLLEAVEAVIELLETLLELNPTNEELLREILKIILRIFELLFELAKKQNDVELLLEAVEAVI<br>ELLETLLELNPTNEELLREILKIILRIFELLFELAKKQNDTELLSEAKEAVAELLETLAELNPTN<br>QELKEEIKKIQERIAELEKELAEKQNA |
| FD33 | MGSSHHHHHHSSGLVPRGSHMENLYFQGSWSGGSGGPEEALEEVEERIEELESALESN<br>PTNEEELREILKKILEIFEELFREAKARNDTELLLEAVEAVIELLETLLELNPTNEELLREILKIIL<br>RIFELLFELAKKLNDTELLLEAVEAVIELLETLLELNPTNEELLREILKIILRIFELLFELAKKLND<br>TELLLEAVEAVIELLETLLELNPTNEELLREILKIILRIFELLFELAKKLNDTELLLEAVEAVIELL<br>ETLLELNPTNEELLREILKIILRIFELLFELAKKLNDTELLSEAKEAVAELLETLAELNPTNQEL<br>KEEIKKIQERIAELEKELAEKLNA |
| FD34 | MGSSHHHHHHSSGLVPRGSHMENLYFQGSWSGGSGGPAEALEEVEERIEELESALASN<br>PTNEEELREILKKILEIFEELFREAKARNDTALLLEAVEAVIELLETLLALNPTNEELLREILKIIL<br>RIFELLFELAKKQNDTALLLEAVEAVIELLETLLALNPTNEELLREILKIILRIFELLFELAKKQN<br>DTALLLEAVEAVIELLETLLALNPTNEELLREILKIILRIFELLFELAKKQNDTALLLEAVEAVIEL<br>LETLLALNPTNEELLREILKIILRIFELLFELAKKQNDTALLSEAKEAVAELLETLAALNPTNQE<br>LKEEIKKIQERIAELEKELAEKQNA |
| FD35 | MGSSHHHHHHSSGLVPRGSHMENLYFQGSWSGGSGGKEEALREVRERIEELKRQLESN<br>PHNEEALREILKKILEIFEELFREAKERNDTELLLRAVRAVIELLKTLLELNPHNEELLREILKII<br>LRIFELLFELAKEQNDTELLLRAVRAVIELLKTLLELNPHNEELLREILKIILRIFELLFELAKEQ<br>NDTELLLRAVRAVIELLKTLLELNPHNEELLREILKIILRIFELLFELAKEQNDTELLLRAVRAVI<br>ELLKTLLELNPHNEELLREILKIILRIFELLFELAKEQNDVELLKRAKEAVDELLRTLAELNPH<br>NEELKEEIKKIEERIEELEKELAEEQND |
| FD36 | MGSSHHHHHHSSGLVPRGSHMENLYFQGSWSGGSGGPEEALAEVEERIAELESALASN<br>PHNEEELREILKKILEIFEELFREAKERNDTELLLAAVEAVIALLETLLALNPHNEELLREILKII<br>LRIFELLFELAKELNDTELLLAAVEAVIALLETLLALNPHNEELLREILKIILRIFELLFELAKELN<br>DTELLLAAVEAVIALLETLLALNPHNEELLREILKIILRIFELLFELAKELNDTELLLAAVEAVIAL<br>LETLLALNPHNEELLREILKIILRIFELLFELAKELNDSELLSAAKEAVDALLETLAALNPNNEE<br>LKEEIKKIEERIEELEKELAEELNS |
| FD37 | MGSSHHHHHHSSGLVPRGSHMENLYFQGSWSGGSGGPEEALEEVEERIEELESALESN<br>PHNEEELREILKKILEIFEELFREAKERNDTELLLEAVEAVIELLETLLELNPHNEELLREILKII<br>LRIFELLFELAKELNDTELLLEAVEAVIELLETLLELNPHNEELLREILKIILRIFELLFELAKELN<br>DTELLLEAVEAVIELLETLLELNPHNEELLREILKIILRIFELLFELAKELNDTELLLEAVEAVIEL<br>LETLLELNPHNEELLREILKIILRIFELLFELAKELNDSELLSEAKEAVDELLETLAELNPNNEE<br>LKEEIKKIEERIEELEKELAEELNS |

| | |
|---|---|
| FD38 | MGSSHHHHHHSSGLVPRGSHMENLYFQGSWSGGSGGPAEALEEVEERIEELESALASNPHNEEELREILKKILEIFEELFREAKERNDTALLLEAVEAVIELLETLLALNPHNEELLREILKIILRIFELLFELAKEQNDTALLLEAVEAVIELLETLLALNPHNEELLREILKIILRIFELLFELAKEQNDTALLLEAVEAVIELLETLLALNPHNEELLREILKIILRIFELLFELAKEQNDTALLLEAVEAVIELLETLLALNPHNEELLREILKIILRIFELLFELAKEQNDSALLSEAKEAVDELLETLAALNPNNEELKEEIKKIEERIEELEKELAEEQNS |
| FD39 | MGSSHHHHHHSSGLVPRGSHMENLYFQGSGGSGGKEKEEALKEILKALEEGGISGELLRQLEEVLKRFLKKAGASEEAILALLEILAALLEGGISIELLLQLLEVLIRFLKKAGLSEEAILALLEILAALLEGGISIELLLQLLEVLIRFLKKAGTSEEEILALLEELAALLEGGISIEELLKRLEEEIKHLKKAGSSW |
| FD40 | MGSSHHHHHHSSGLVPRGSHMENLYFQGSGGSGGPSIEAALKELEEALEAALRELLEAAGVDAEAIERVLEIILRALEEILEIARRAGDDPSFLLALFELAIALLLALFELLVAAGVDAEAIARVLLIILRALIEILEIAVRAGDDPSFLLALFELAIALLLALFELLAAAGVDAEAIARVLLIILRALIEILEIAVRAGDDPSELLALFEAAIARLLALFEALLAQGVDAEAIARRLLEELRKLIERLEEEVRASDEW |

Table S2: Biochemical and structural properties of the proteins tested in this study.

| Name | Size (nm) (l x w x h) | Repeat Spacing (nm) | Net Charge | Number of carboxylate groups | Number of carboxylates at the interface | Ratio of aspartate to glutamate | Solvent accessible surface area ($nm^2$) | Percent surface hydrophobic |
|---|---|---|---|---|---|---|---|---|
| BSA | 8.3 x 7.4 x 6.1 | n.a. | -18 | 98 | n.a. | 39/59 | 279.33 | 57.8 |
| FD15 | 6.4 x 5.5 x 1.8 | 0.87 | -32 | 67 | 30 | 10/57 | 154.54 | 51.5 |
| DHR49-Neg | 6.7 x 3.7 x 2.3 | 1.01 | -54 | 72 | 42 | 30/42 | 115.39 | 46.4 |
| FD31 | 7.9 x 4.6 x 2.4 | 1.12 | -47 | 81 | 36 | 5/76 | 150.93 | 49.3 |
| FD31-Rep3 | 4.6 x 4.6 x 2.4 | 1.12 | -26 | 45 | 18 | 2/43 | 89.93 | 50.1 |
| FD31-Rep9 | 11.2 x 4.6 x 2.4 | 1.12 | -68 | 117 | 54 | 8/109 | 220.13 | 47.4 |
| FD31-Gln-Chkr | 7.9 x 4.6 x 2.4 | 1.12 | -29 | 63 | 18 | 5/58 | 152.12 | 48.9 |
| FD31-Asp | 7.9 x 4.6 x 2.4 | 1.12 | -47 | 81 | 36 | 29/52 | 148.77 | 48.7 |
| FD31-Lys-Chkr | 7.9 x 4.6 x 2.4 | 1.12 | -23 | 69 | 24 | 5/64 | 155.38 | 50.2 |

### Supporting Figures

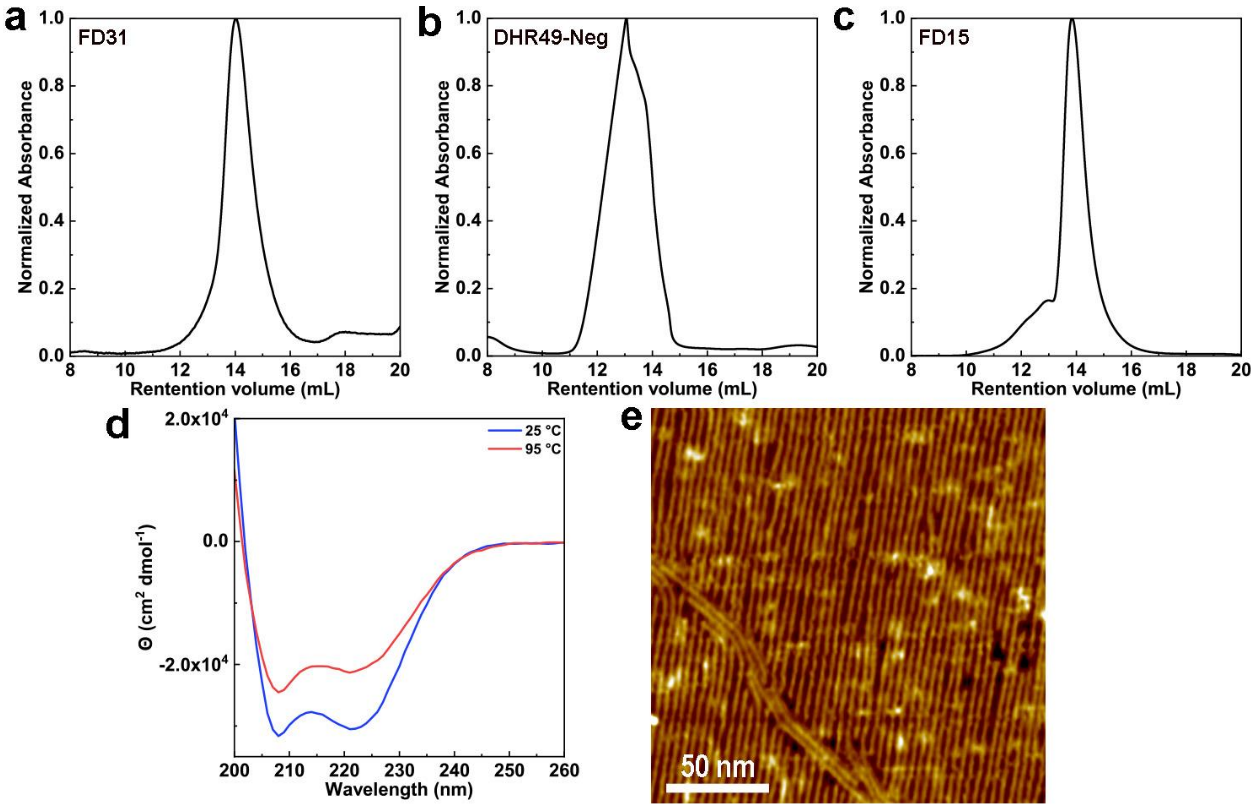


Fig. S1. Biophysical characterization of selected proteins. Normalized ultraviolet absorbance (A230) of a size exclusion chromatography trace for (a) FD31, (b) DHR49-Neg, (c) FD15. (d) Circular dichroism scans from at 25 °C (blue) and 95 °C (red) for DHR49-Neg. (e) AFM image of DHR49-Neg assembly on mica in 3 M KCl solvent showing fiber-like behavior driven by end-to-end oligomerization.

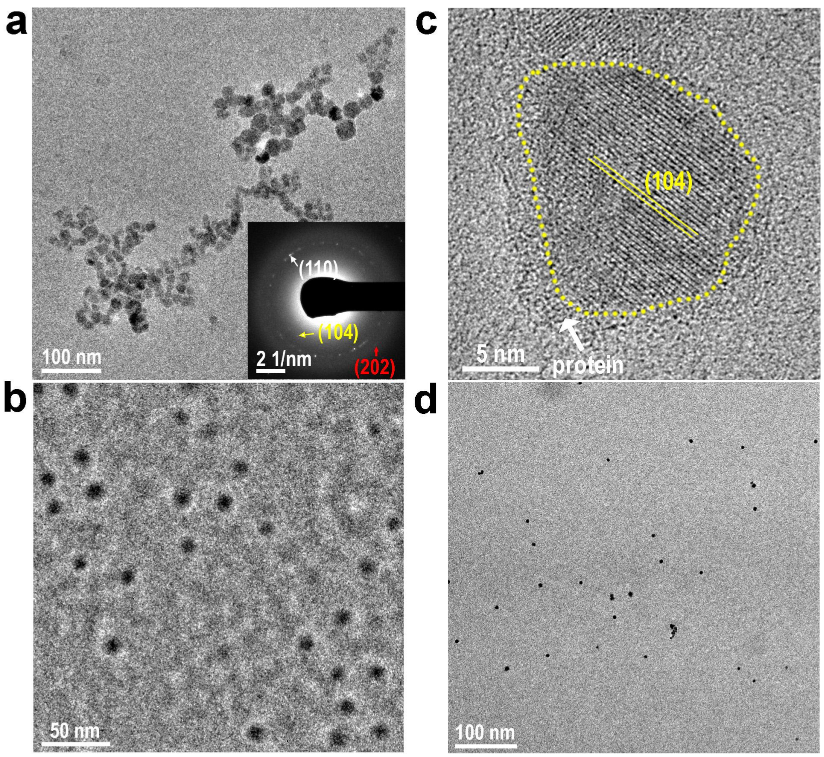


**Fig. S2.** TEM images of calcite nanocrystals in the presence of 1 μM FD31. (a) TEM and corresponding SAED images confirm the nanocrystals are calcite. (b) Cryo-TEM image of $CaCO_3$ particles at 5 mins. (c) HR-TEM image of calcite nanocrystals. The amorphous objectives with low contrast are suggestive of a shell around the nanocrystals, which could be absorbed protein. (d) LP-TEM image shows ~ 5 nm CaCO3 nanoparticles formed in the mineralization solution.

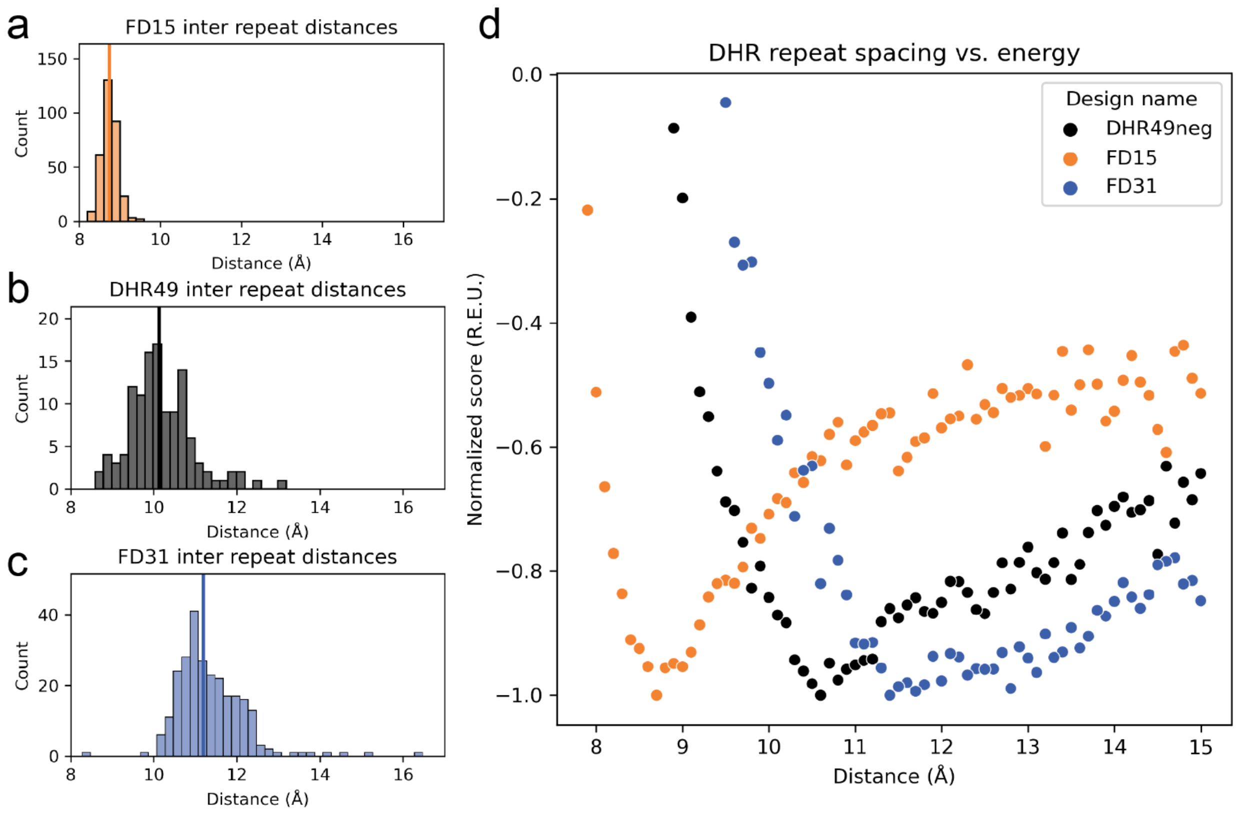


Fig. S3. Distribution of distances between α-carbon atoms in adjacent repeats within (a) FD15 crystal structure, (b) the DHR49-Neg crystal structure (pdb id: 5CWJ) that DHR49-Neg was derived from, and (c) the AlphaFold model of FD31. Lines show median values for each protein (FD15=8.7Å; DHR49-Neg=10.1Å; FD31=11.2Å). (d) Normalized predicted protein energy in Rosetta energy units (R.E.U.) vs. repeat-repeat distance (Å) for the lowest energy DHR models constrained to flat conformations with a specified repeat-repeat distance.

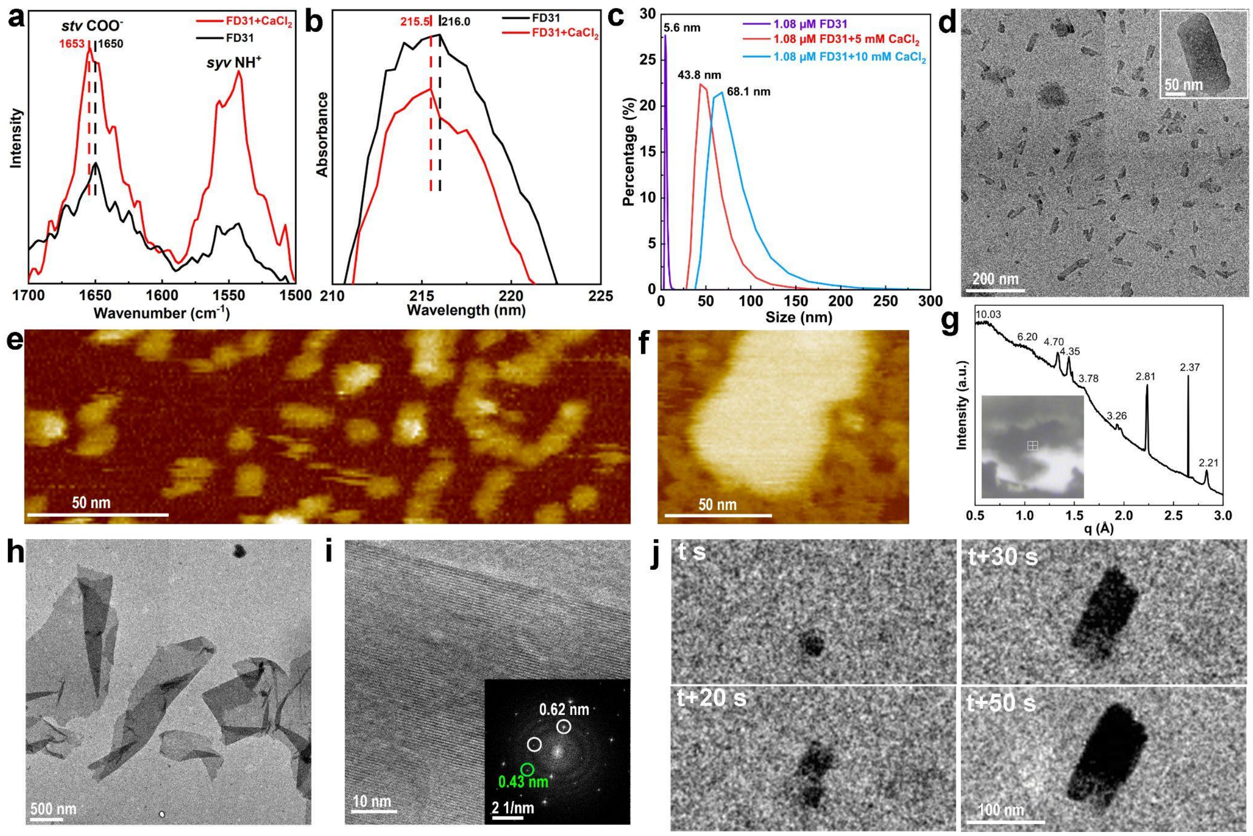


Fig. S4. Structural characterization of the FD31-Ca complex. (a) Liquid-phase ATR-FTIR spectra and (b) UV-vis spectra of pure FD31 (black) and FD31 plus $CaCl_2$ (red) solutions. The peak shift shows the coordination of Ca with carboxylate groups on the protein. (c) DLS size measurements of FD31 with different $CaCl_2$ concentrations (0, 5, and 10 mM). (d) LP-TEM image showing the sheet-like assemblies. (e-f) AFM images showing the rod-like individual FD31 molecules and FD31-Ca complex on mica with CaCl2. Panel e reveals a rod-like shape of the FD31 protein molecules dispersed in MOPS buffer solution with a size of ~8 nm by 5 nm by 2 nm (Length by width by height), which is consistent with their simulated “Flattened” Rosetta structure model (~7.8 nm by 4.5 nm by 2.5 nm). (g) Beamline XRD of assembled protein structure. (h-i) Representative TEM image showing the sheet-like assemblies (j) Time-dependent LP-TEM images show the formation of the FD31-Ca complex.

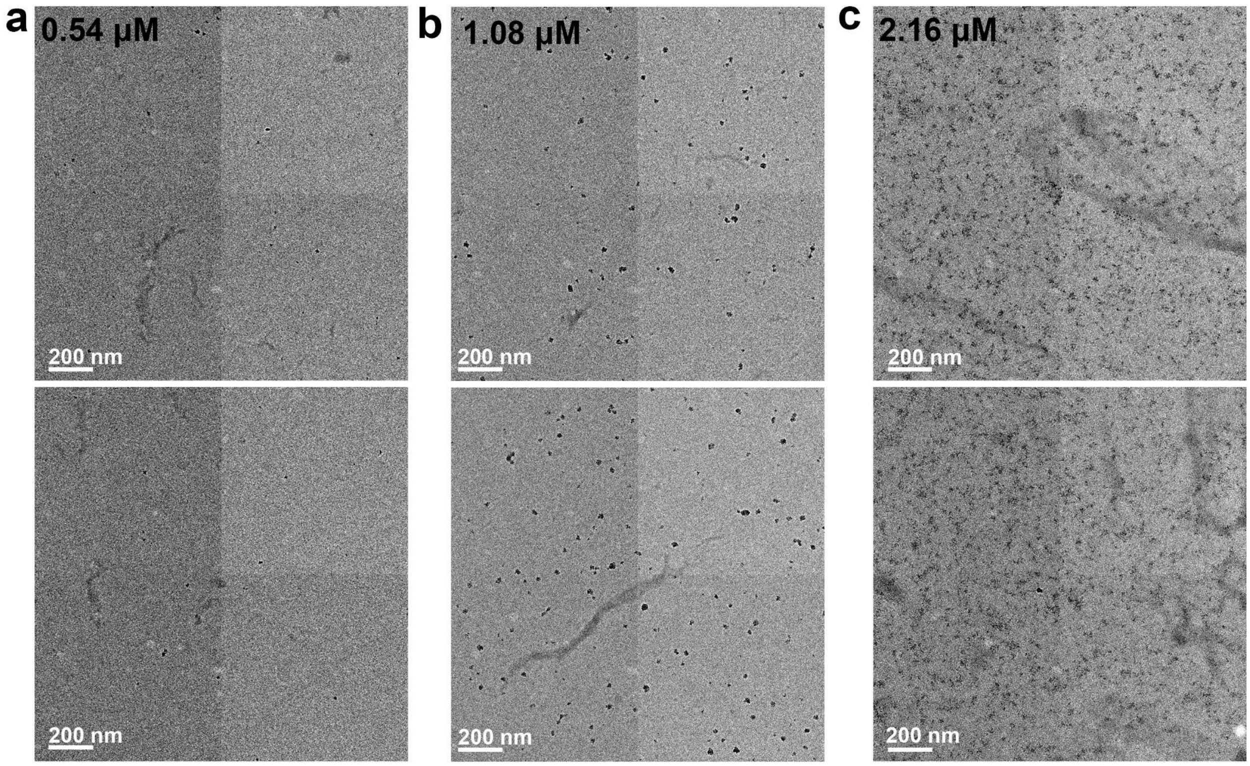


**Fig. S5.** FD31 concentration effects on $CaCO_3$ nucleation in supersaturated solutions containing 5 mM $CaCl_2$ and 5 mM $NaHCO_3$. FD31 was incubated with 10 mM $CaCl_2$ for 5 minutes prior to addition of an equal volume of 10 mM $NaHCO_3$. TEM samples were prepared 10-20 minutes after addition of $NaHCO_3$. (a-c) Representative TEM images show the resulting $CaCO_3$ particles on copper grids in the presence of 0.54, 1.08, 2.16 μM FD31 proteins.

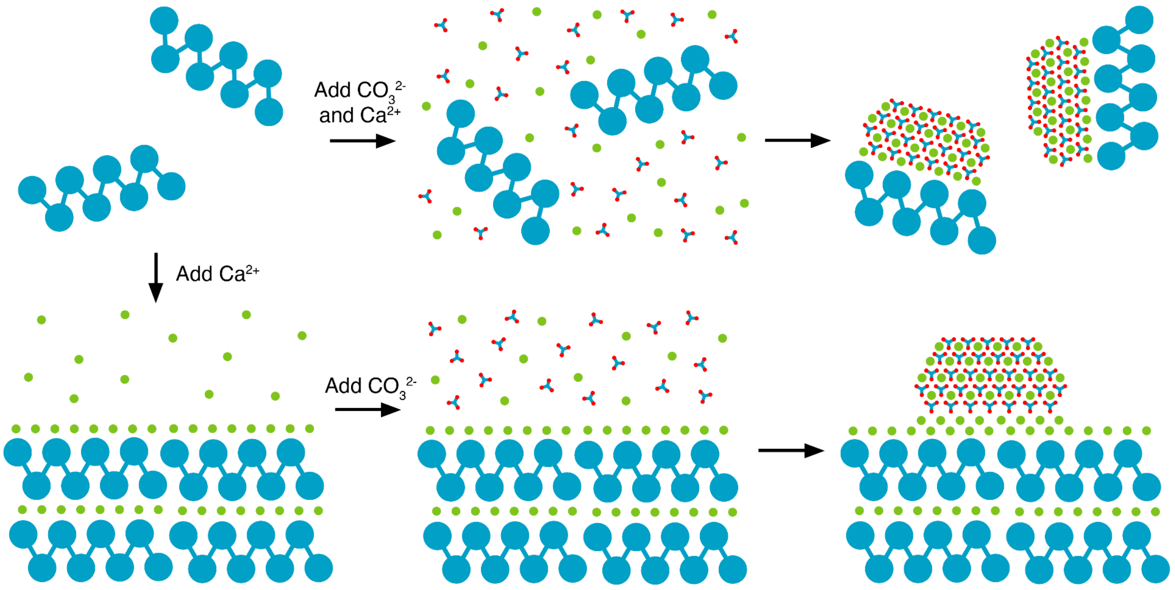


Fig. S6. Proposed nucleation pathways for the FD31 system. (Top) when $Ca^{2+}$ and $CO_3^{2-}$ are mixed with protein simultaneously, nucleation of calcite is driven by protein monomers. (Bottom) when $Ca^{2+}$ is added first, protein-$Ca^{2+}$ assemblies form and serve as templates for calcite upon addition of $CO_3^{2-}$.

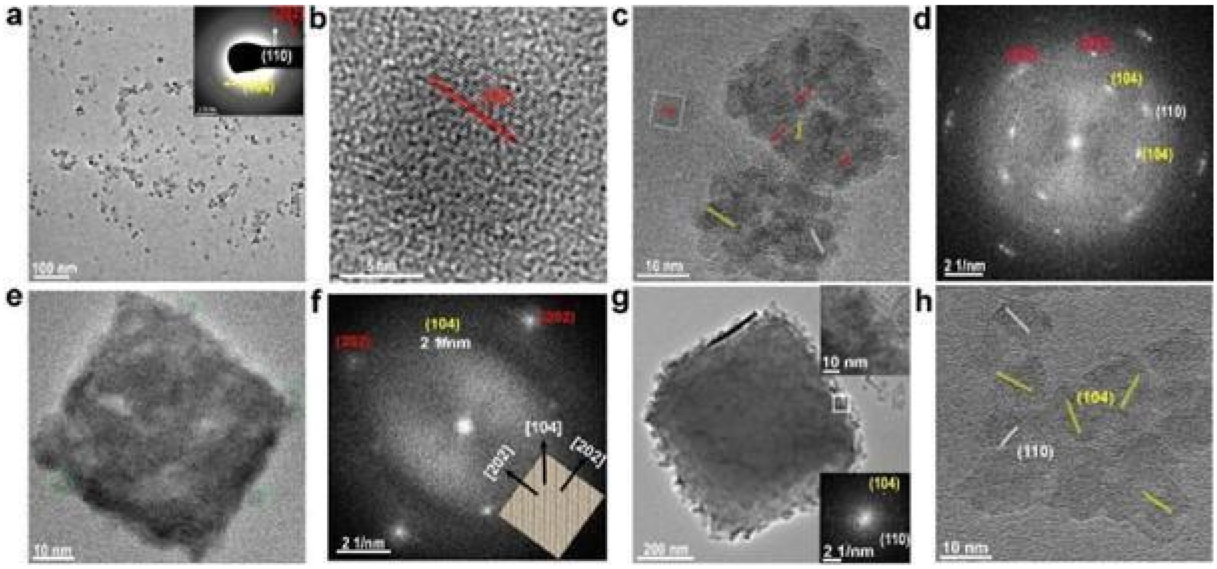

**Fig. S7.** The $CaCO_3$ crystallization process in the presence of DHR49-Neg. (a) TEM and inserted SAED images show that the nanocrystals are calcite. (b) HR-TEM image shows an individual nanocrystal with (202) lattice direction. (c, d) HR-TEM and corresponding FFT images show aggregated calcite single crystals. (e, f) HR-TEM and corresponding FFT images show a cubic-like single crystal with exposed (202) facets. (g, h) Individual rhombohedral calcite with some nanocrystals on or around it. Inserted HR-TEM and FFT images in panel “g” to confirm that calcite nanocrystals are incorporated into the pre-existed calcite.

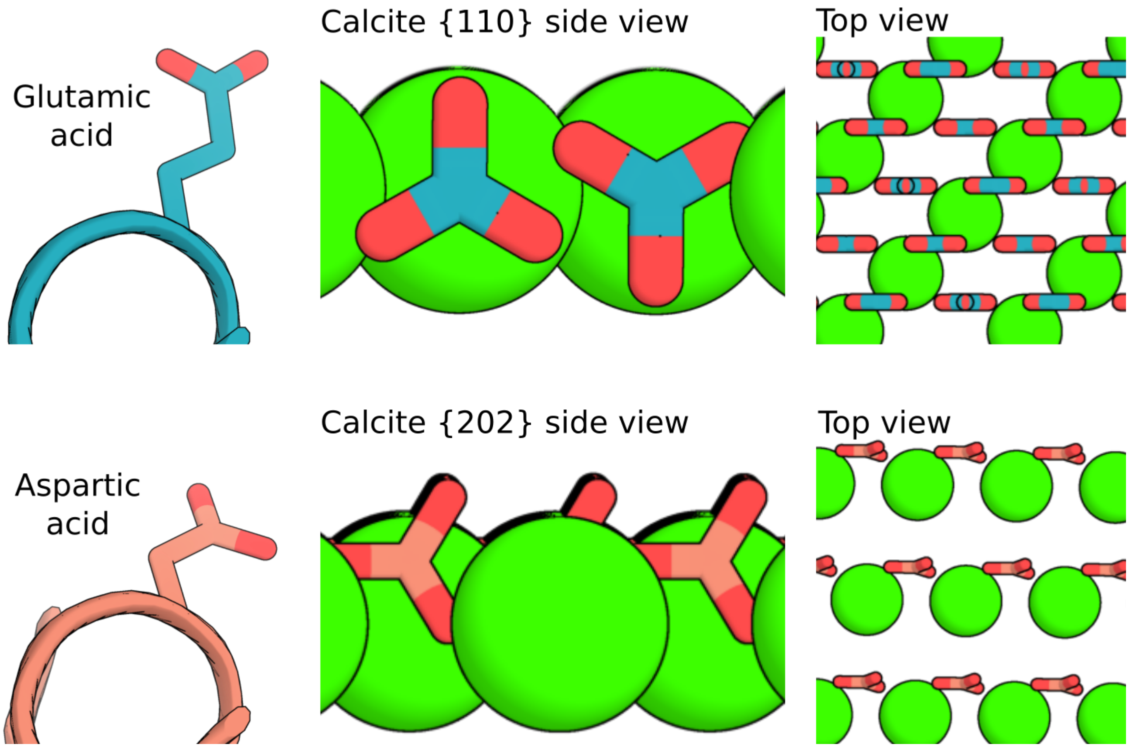


**Fig. S8.** Orientations of the carboxylate groups in different binding moieties could explain the bias towards the formation of a particular calcite facet.

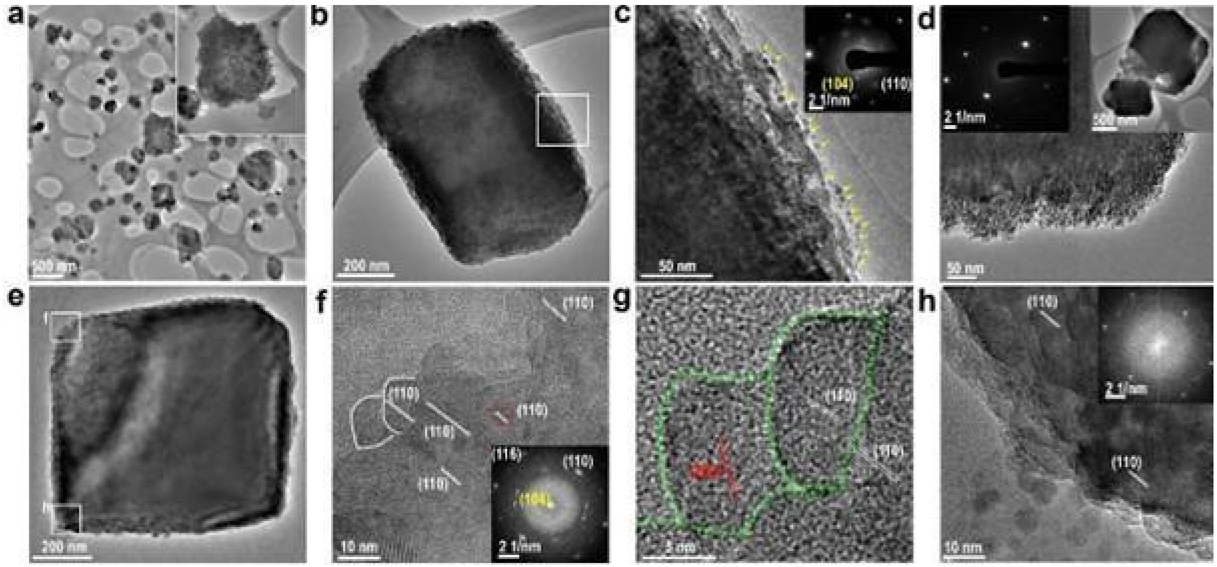


**Fig. S9.** TEM images of calcite at 30 mins in the presence of 1 μM FD31 protein. (a) Irregular calcite consisting of nanocrystals. Inset shows an individual calcite aggregate. (b, c) One calcite crystal is surrounded by some calcite nanoparticles. Inserted SAED image confirms the single crystalline nature. (d) Rhombohedral calcite crystals consist of smaller nanocrystals. (e) TEM image shows a rhombohedral calcite with a size of ~800 nm. (f,g,h) Higher magnification views of the corners of the particle in (e). (f) HR-TEM image of the corner region showing the same (110) lattice orientation as in the crystal body. (g) HR-TEM image showing several single-crystal units on the surface. (h) HR-TEM image showing numerous small nanocrystals located around the rhombohedral calcite.

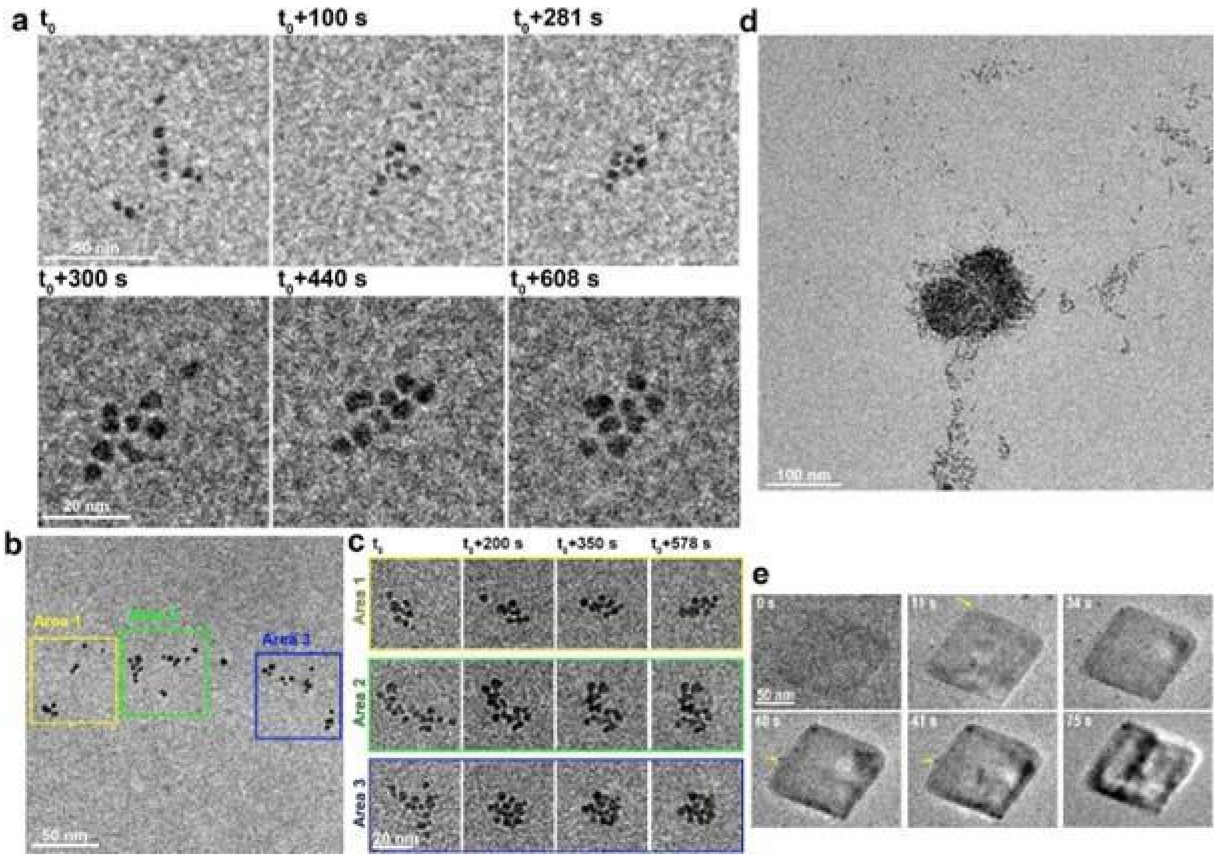


**Fig. S10.** LP-TEM observation of dynamical behavior of calcite nanoparticles in liquid-cell. (a) Sequential TEM images showing the aggregation process of multiple particles. (b) LP-TEM image showing the initial distribution of particles. (c) Series of TEM images showing the formation of three aggregates. (d) LP-TEM image showing the aggregated particles. (e) Sequential TEM images show the attachment of particles into pre-existed rhombohedral calcite.

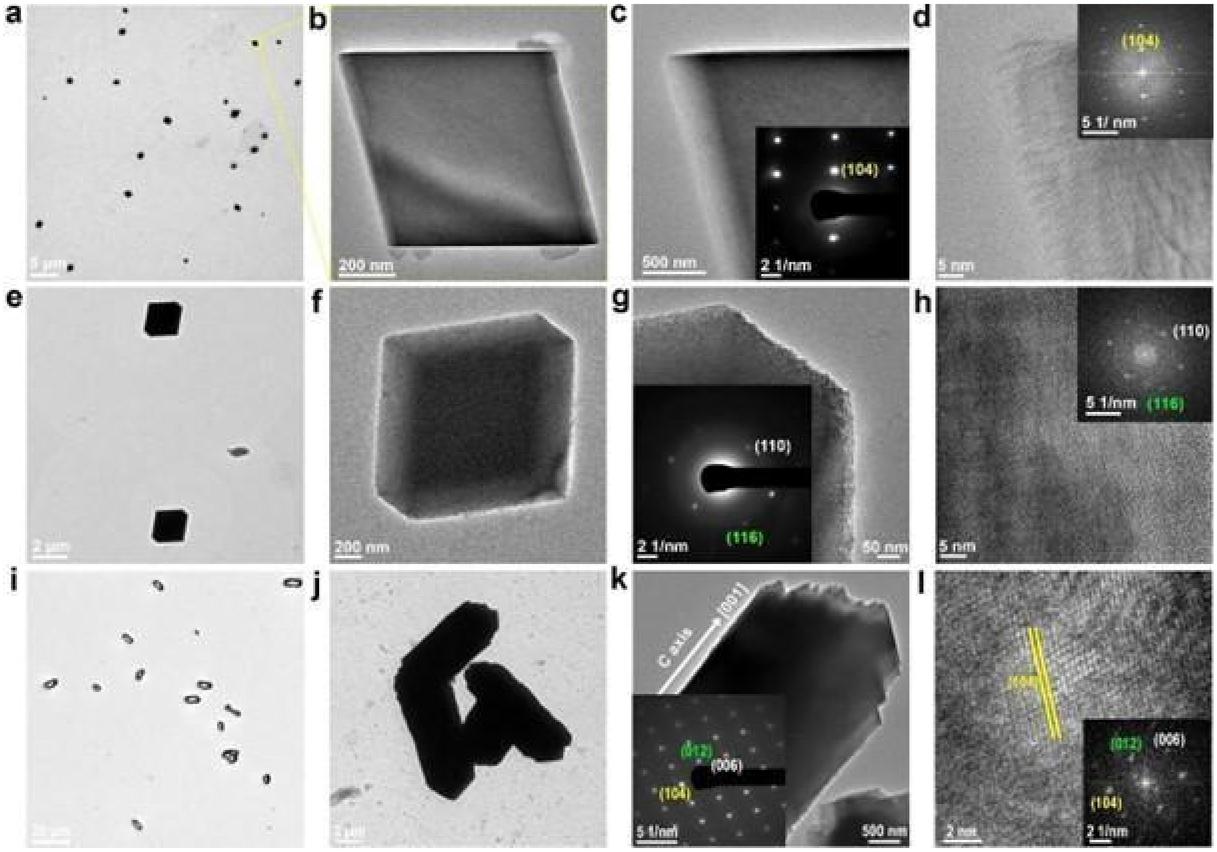


**Fig. S11.** The resulting $CaCO_3$ morphology after 8 hours without additives or with proteins of different concentrations in supersaturated solutions containing 5 mM CaCl2 and 5 mM $NaHCO_3$. (a-d) TEM and SAED images show the formation of perfect rhombohedral calcite without the involvement of proteins. (e-h) TEM and SAED images show the formation of rough rhombohedral calcite with the addition of 1.08 μM FD31 proteins. (i) OM (k), TEM and SAED images show the formation of rod-like calcite with the addition of 4.32 μM FD31 protein.

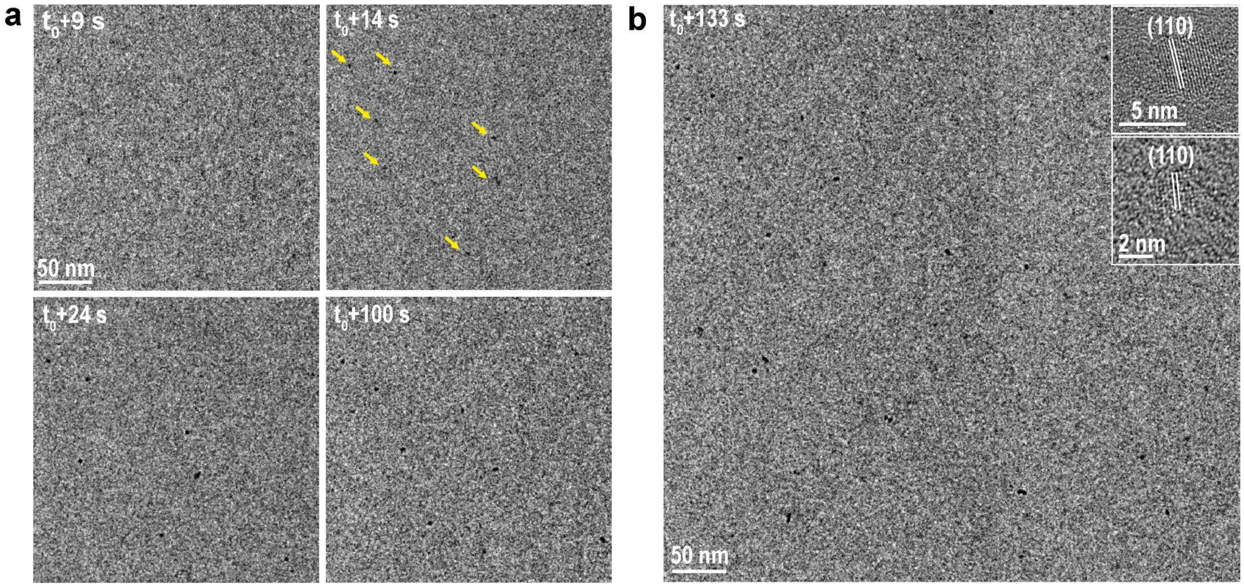


**Fig. S12.** In situ LP-TEM observation of $CaCO_3$ crystallization process in the presence of FD31-Rep9 protein. (a) Sequential TEM images showing the nucleation of CaCO3 nanoparticles. (b) TEM image showing multiple $CaCO_3$ nanoparticles in the liquid cell. Inserted HR-TEM image confirms the nanocrystals are calcite.

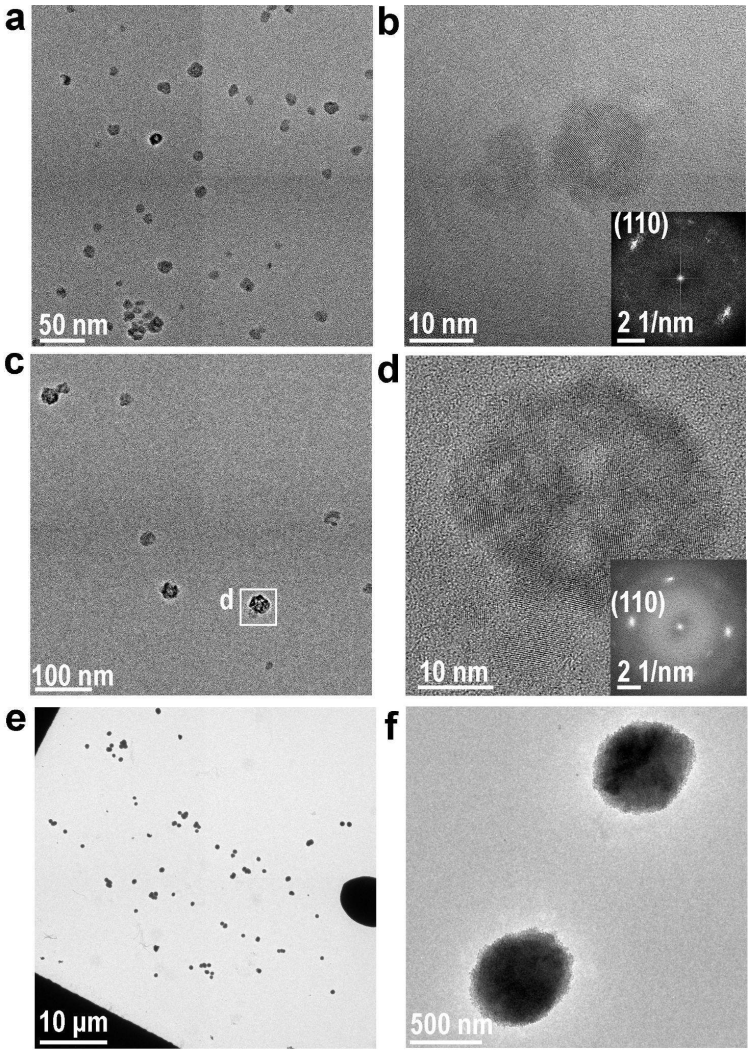


**Fig. S13.** Resulting $CaCO_3$ in the presence of different FD31-mutants. (a-b) TEM and SAED images show the formation of calcite-dominant nanocrystals in the presence of FD31-Gln-Checker. (c-d) TEM and FFT images show some calcite nanocrystals in the presence of FD31-Asp. (e-f) TEM and SAED images show the formation of a dominant vaterite phase in the presence of FD31-Lys-Checker.

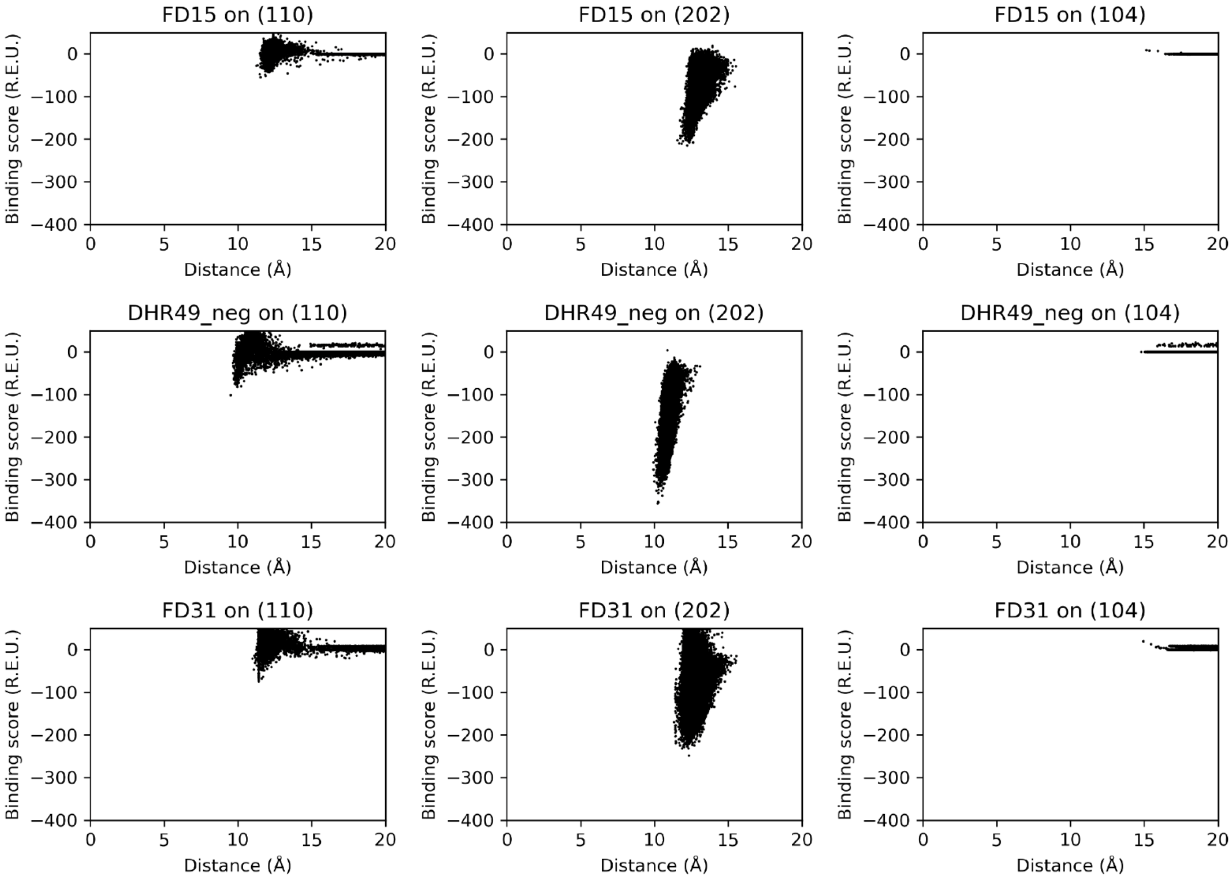


Fig. S14. Binding score vs. distance from the protein's center of mass to the surface in all Rosetta docking trajectories of DHRs on calcite surfaces. To allow for the possibility of small conformational changes in the proteins upon adsorption to the mineral surface, idealized models of each DHR with a range of inter-repeat spacings at 0.1Å increments were docked on the surfaces (8.0Å-9.7Å for FD15, 9.6Å-11.6Å for DHR49-Neg, and 10.4Å-12.4Å for FD31). All three proteins showed binding signals for the (110) and (202) calcite surfaces and not for the (104) surface.

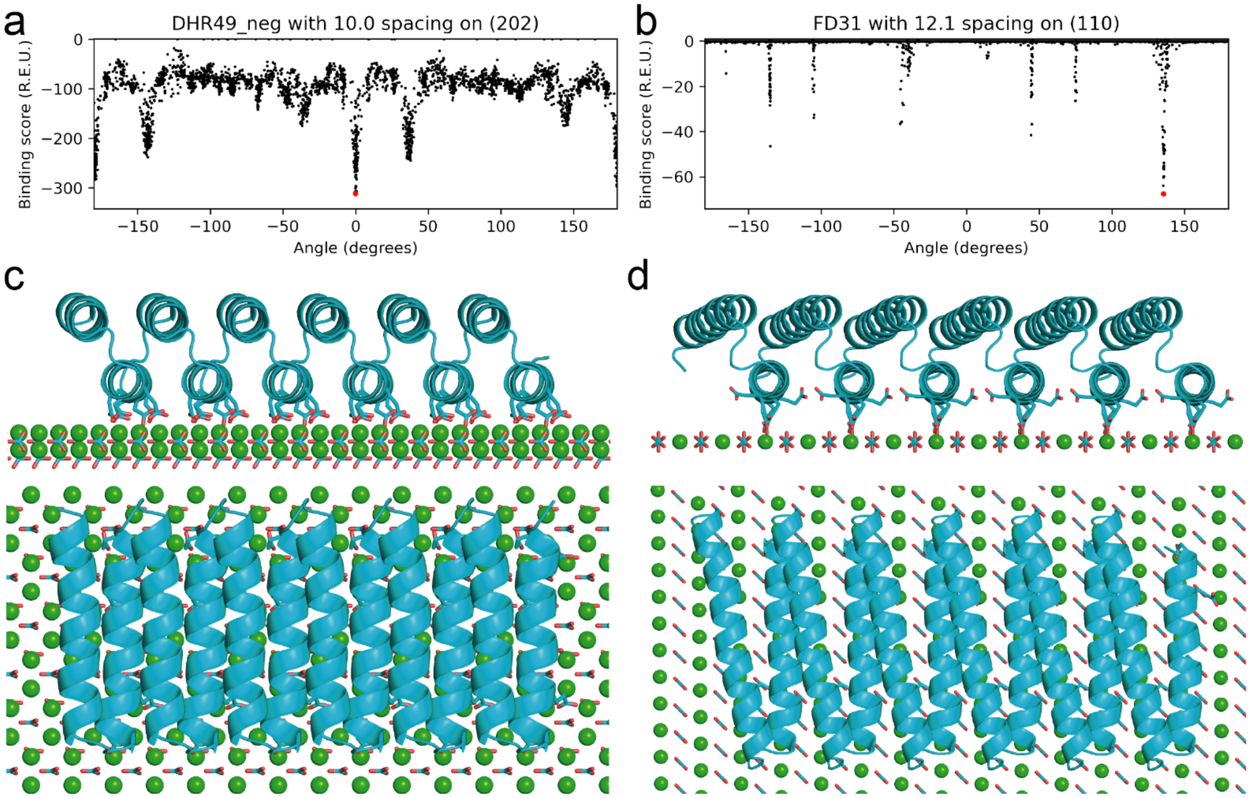


Fig. S15. Rosetta binding scores vs. angle of DHRs adsorbed on calcite surfaces and illustrations of how these proteins may form lattice matches with the surfaces. The angles are measured between the projection of the protein's repeat axis onto the plane of the calcite surface and an arbitrary reference vector that is parallel to the surface. (a) The binding scores vs. angle of DHR49-Neg idealized with a 10.0 Å repeat spacing on the (202) surface and (b) of FD31 idealized with 12.1 spacing on the (110) surface. (c) Lowest energy structure of the DHR49-neg model on (202), indicated by the red dot in panel (a). (d) Side and top views of the lowest energy structure of the FD31 model on (110), indicated by a red dot in panel (b).

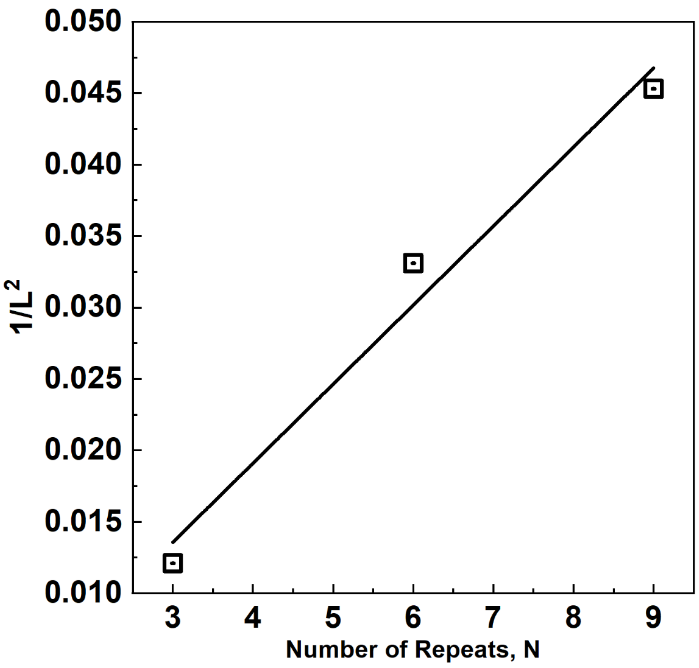


Fig. S16. Dependence of nanocrystal size on length of DHR protein template, where L is the average nanocrystal diameter and N is the number of repeats in the protein. The nanocrystal surface area is proportional to $L^2$ and the protein surface area is proportional to N.
